# Experimental evolution reveals a general role for the methyltransferase Hmt1 in noise buffering

**DOI:** 10.1101/714949

**Authors:** Shu-Ting You, Yu-Ting Jhou, Cheng-Fu Kao, Jun-Yi Leu

## Abstract

Cell-to-cell heterogeneity within an isogenic population has been observed in prokaryotic and eukaryotic cells. Such heterogeneity often manifests at the level of individual protein abundance and may have evolutionary benefits, especially for organisms in fluctuating environments. Although general features and the origins of cellular noise have been revealed, details of the molecular pathways underlying noise regulation remain elusive. Here, we used experimental evolution of *Saccharomyces cerevisiae* to select for mutations that increase reporter protein noise. By combining bulk segregant analysis and CRISPR/Cas9-based reconstitution, we identified the methyltransferase Hmt1 as a general regulator of noise buffering. Hmt1 methylation activity is critical for the evolved phenotype, and noise buffering is primarily achieved via two Hmt1 methylation targets. Hmt1 functions as an environmental sensor to adjust noise levels in response to environmental cues. Moreover, Hmt1-mediated noise buffering is conserved in an evolutionarily distant yeast species, suggesting broad significance of noise regulation.

**Author Summary:** Cell-to-cell heterogeneity within an isogenic population has been observed in prokaryotic and eukaryotic cells. Such heterogeneity often manifests at the level of individual protein abundance and may have evolutionary benefits, especially for organisms in fluctuating environments. Here, we used experimental evolution of *Saccharomyces cerevisiae* to select for mutations that increase reporter protein noise and identified the methyltransferase Hmt1 as a general regulator of noise buffering. Hmt1 is a central hub protein that is involved in multiple basic cellular pathways, including chromatin remodeling/transcription, translation, ribosome biogenesis, and post-transcriptional regulation. Our results show that Hmt1 constrains the noise level of multiple cellular pathways under normal conditions, so the physiology of individual cells in a population will not deviate too much from optimal peak fitness. However, when cells encounter environmental stresses, HMT1 is quickly down-regulated and expression noise is enhanced to increase the likelihood of population survival. Moreover, the noise buffering function of Hmt1 is conserved in *Schizosaccharomyces pombe* that diverged from the common ancestor of *Saccharomyces cerevisiae* more than 400 million years ago. Since the Hmt1 network is conserved from yeast cells to human, it is quite possible that Hmt1-mediated noise buffering also operates in multicellular organisms.

## Introduction

Genetically identical cells grown under homogeneous conditions can still exhibit heterogeneous phenotypes. This heterogeneity is ubiquitous and manifests at different levels, from individual protein concentrations [1] to cell physiology [2, 3]. Although phenotypic heterogeneity only exists transiently, it can lead to deterministic outcomes. In multicellular organisms, a stochastic difference in the initial cell state can result in different cell fates during development [4, 5]. Moreover, stochastic variation in gene expression has been shown to determine the outcome of inherited detrimental mutations [6, 7], representing a possible cause for the incomplete penetrance observed in many human diseases. In microbial cells, levels of pre-existing heterogeneity can influence population fitness upon exposure to unpredictable environmental change [8, 9]. This “bet-hedging strategy” is commonly used by microorganisms to ensure population survival without the fitness cost of developing complex regulatory networks that respond to fluctuating environments [10].

At the gene expression level, pre-existing cell-to-cell heterogeneity (or “cellular noise”) mainly originates from the stochasticity inherent to molecular processes (such as transcription factor binding to target sequences) and fluctuating levels or activities of factors critical to those processes (such as RNA polymerase II or ribosomes) [1, 11]. Genome-wide studies have shown that low-abundance proteins often present higher expression noise, which is consistent with the greater variability of infrequent events [12, 13].

However, some studies have revealed certain pathway-specific patterns in noise levels. For example, housekeeping genes tend to have lower noise, whereas environment-responsive genes are often noisier [14, 15], perhaps because fluctuations in housekeeping genes may compromise essential cellular functions and noisy environment-responsive genes can exert a bet-hedging function. The observed patterns in these two types of genes indicate that selection operates on protein noise levels or levels have been adjusted according to potential costs and benefits over the course of evolution. Moreover, a study comparing young and old mice showed that heart cells isolated from old mice exhibit higher gene expression noise than those isolated from young mice [16], suggesting that noise levels are tightly controlled in young healthy cells but the control systems deteriorate with age.

How do cells adjust protein noise? Several general features have been associated with expression noise including network topology, cellular compartmentalization, molecular chaperone abundance, nucleosome occupancy, and promoter architecture [7, 17–19]. Genetic studies have also identified mutations that alter local or general noise levels [5, 11, 20, 21]. Nonetheless, how cells respond to growth conditions and integrate different pathways to fine-tune protein noise remains insufficiently characterized.

To understand how protein noise is regulated, we used experimental evolution of *Saccharomyces cerevisiae* to search for mutations that increased the protein noise of different reporter genes. After 35 cycles of selection, two of the evolved lines (*TDH2-GFP*- and *TYS1-GFP*-carrying lines) exhibited increased noise levels without a concomitant reduction in protein abundance. We show that increased protein noise in the evolved line carrying the *TDH2* reporter gene is not specific to the *TDH2*-related pathway, suggesting that the evolved mutations have a general effect on protein noise regulation. We identified the methyltransferase Hmt1 as the major contributor of the evolved phenotype. Further experiments revealed that noise regulation is mediated by methylation of multiple downstream targets of Hmt1 and that *HMT1* expression is often attenuated under stress conditions. Our results suggest that Hmt1 functions as a master coordinator of bet-hedging strategies in response to environmental stress.

## Results

### Experimental evolution of increased protein noise in budding yeast

A previous study showed that alternating selection between highest- and lowest-expression subpopulations could efficiently enrich promoter variants for high transcriptional noise in bacterial cells [22]. We hypothesized that a similar selection strategy might allow us to “evolve” yeast cells to increase the protein noise of reporter genes (Fig. 1A). Eight genes (*ADK1*, *APA1*, *PCM1*, *RPL4B*, *SAM4*, *TDH2*, *TPD3*, and *TYS1*) selected from distinct cellular pathways were fused with GFP to generate our reporters (see Materials and Methods). Evolving lines carrying individual reporter genes were subjected to alternating selection between the top 5% and bottom 5% of total populations in terms of their GFP intensity. We also treated cells with a mutagen (2.8% ethyl methanesulfonate, EMS) before each selection cycle to increase the genetic diversity of evolving populations. After 35 cycles of selection, half of the evolved lines (including the strains carrying *RPL4B*, *TDH2*, *TPD3*, or *TYS1* reporter genes) exhibited significantly increased reporter noise (Fig. 1B).

**Fig. 1.**
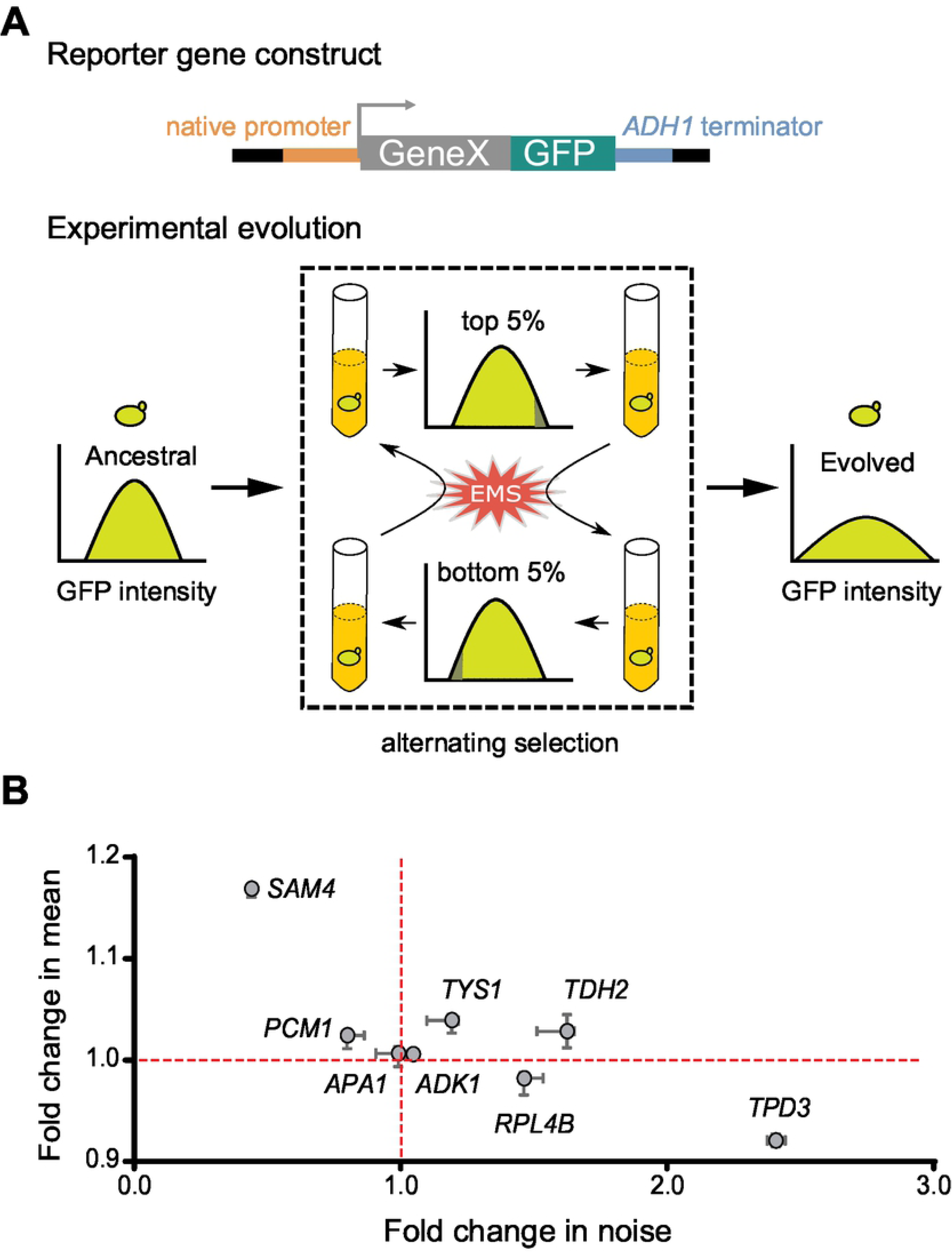
Yeast cells exhibit high noise levels upon experimental evolution. (A) Schematic of our evolution experiment. Cells were treated with 2.8% EMS, regrown for 12 h, and then selected for the top 5% (or bottom 5%) of the population in terms of GFP intensity. Alternating selection enriched for mutant cells that presented higher expression noise in terms of GFP intensity. (B) Most of the evolved clones exhibit increased expression noise. Single clones were isolated from eight evolved cultures and their reporter gene expression was measured. The x- and y-axes represent fold-change in noise and mean expression, respectively, after evolving. The median and range for 4-5 replicates for each evolved clone are indicated by circles and error bars, respectively. The red lines indicate values for the ancestral line.

Among them, the Tdh2-GFP- and Tys1-GFP-carrying lines also presented increased mean protein intensity, ruling out the possibility that the increased noise was due to reduced mean protein signal intensity. These results indicate that our selection regime effectively increased protein noise upon experimental evolution. In order to dissect the genetic basis of noise regulation, we selected the Tdh2-GFP-carrying line for further analysis since it exhibited the greatest increase in noise without a concomitant decrease in mean signal intensity.

### Isogenic cells with high and low Tdh2 levels have fitness advantages under different conditions

Bet-hedging is a commonly adopted survival strategy among microorganisms for spreading the risk of encountering hostile environments [9, 23]. Tdh2 protein is a glyceraldehyde-3-phosphate dehydrogenase involved in glycolysis/gluconeogenesis and it has been shown to help cells resist oxidative stress during the stationary phase [24]. We tested if cells with high or low Tdh2-GFP levels represent different physiological states and if they exhibit fitness advantages under different conditions. To do this, we isolated individual stationary phase cells presenting different levels of Tdh2-GFP using a cell sorter and examined their phenotypes.

Consistent with a previous observation [24], upon H_2_O_2_ treatment, cells with high Tdh2-GFP levels had higher survival rates than those with low Tdh2-GFP (Fig. 2A). Interestingly, when we provided fresh nutrients, the cells with high levels of Tdh2-GFP tended to re-enter the cell cycle more quickly than those with low Tdh2-GFP levels, despite cells with different levels of Tdh2-GFP exhibiting no difference in survival rates under this condition (Fig. 2B). This variation in cell cycle re-entry is reminiscent of the divergent germination times observed in same populations of plants or fungi, which has been suggested to be a risk-spreading strategy to enhance long-term survival [25–27]. Divergent cell cycle re-entry times can prevent an entire cell population from going extinct upon occurrence of an unpredicted environmental catastrophe. To test this hypothesis, we collected cells displaying either high or low Tdh2-GFP and challenged the cells with heat stress either before or after the cells had been re-fed with fresh nutrients. We found that cells with low Tdh2-GFP had a survival rate 3-fold greater than that of cells with high Tdh2-GFP upon encountering heat stress after nutrient refreshment. Survival rates were similar and independent of Tdh2-GFP levels for cells either non-stressed or stressed before nutrient refreshment and (Fig. 2C).

**Fig. 2.**
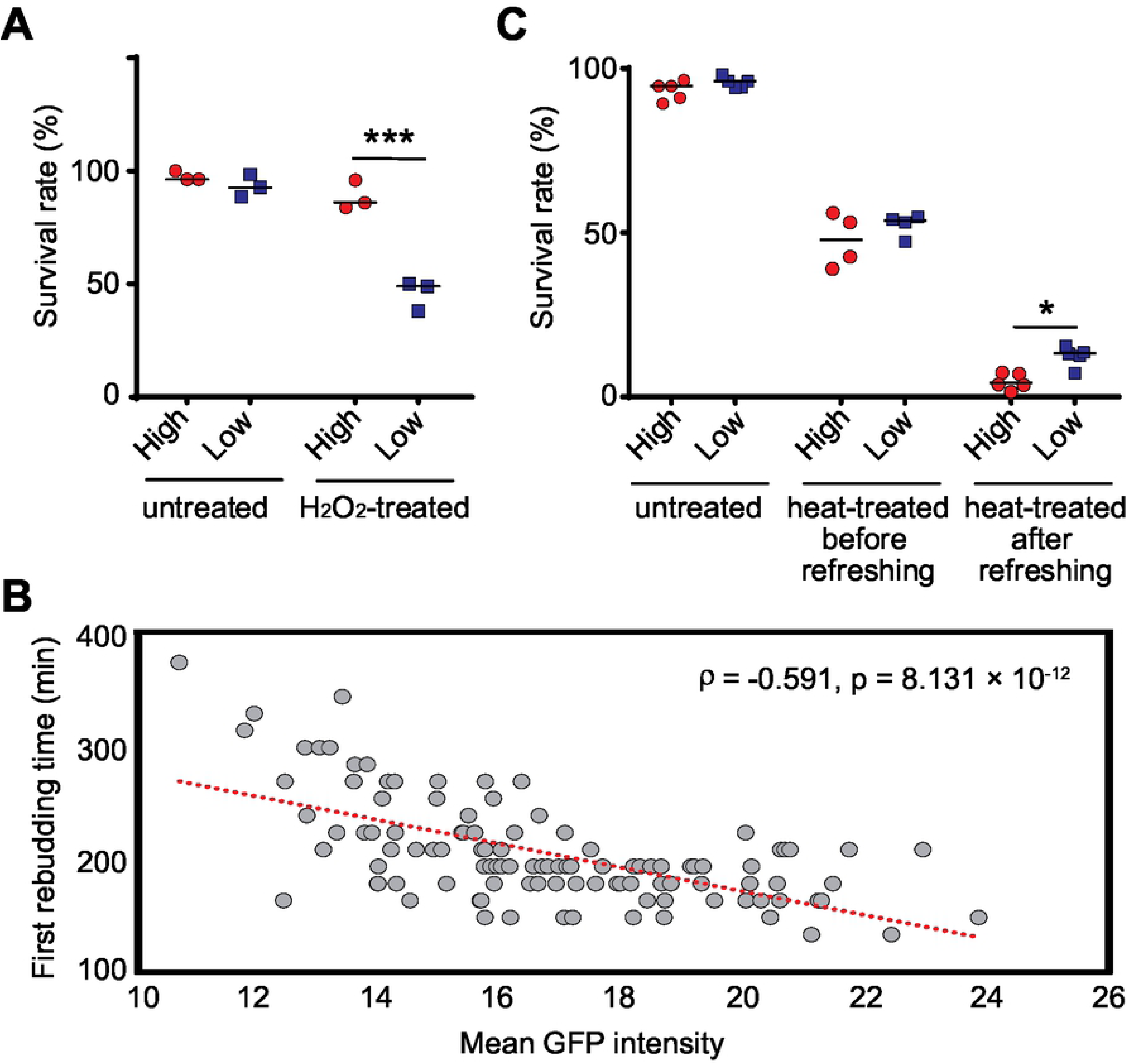
Cells with different levels of Tdh2-GFP exhibit different physiological states. (A) Stationary-phase cells with high Tdh2-GFP levels (red circles) survive better than those with low levels (blue squares) after growing in H_2_O_2_-containing medium (Fisher’s exact test, *p* = 0.1099 for untreated samples, *p* = 1.127 × 10^-4^ for H_2_O_2_-treated samples). Single cells with high or low GFP intensities were sorted and plated on the same plates with or without 4.4 mM H_2_O_2_. Survival rates were determined by counting colony-forming units after 5 days of growth. (B) Stationary-phase cells with high Tdh2-GFP tend to re-enter the cell cycle faster than those with low Tdh2-GFP signal. Each dot of the scatter plot represents data from a single cell. Unsorted stationary-phase cells were placed on YPD agarose pads and only unbudded cells were monitored using time-lapse microscopy. The x-axis indicates the initial Tdh2-GFP signal intensity for each cell and the y-axis indicates the first rebudding time. The red dotted line represents a linear regression (Spearman’s rank correlation, n=111, *p* = 8.131 × 10^-12^). (C) Stationary-phase cells with low Tdh2-GFP signal survive better than those with high signal when cells encounter heat stress after being re-fed with fresh nutrients (one-sided Wilcoxon rank-sum test, n=4-5; *p* = 0.1995 for untreated samples, *p* = 0.2426 for heat-treated samples before nutrient refreshment, *p* = 0.0159 for heat-treated samples after nutrient refreshment). Survival rates were determined by counting colony-forming units after 3 days of growth. The median value for replicates is indicated by solid horizontal lines among groups of data points. \**p* < 0.05, \*\*\**p* < 0.001.

We also found that high and low levels of Tdh2-GFP represent transient states, but not genetic modifications, of the cells. When we propagated cells sorted into high and low Tdh2-GFP populations, Tdh2-GFP intensities of both populations reverted to a level similar to that of the initial unsorted population after a few generations (Fig. S1). Together, our results demonstrate that having cells with different levels of Tdh2 in an isogenic population is advantageous in different environments, revealing the risk-spreading benefit of expression noise.

### Increased noise is not limited to the *TDH2*-related pathway in the Tdh2-GFP evolved line

Before characterizing the detailed phenotypes of the evolved Tdh2-GFP-carrying line, we examined the cell populations and found that the signal for increased noise was not bimodal (Fig. S2A). We further confirmed that Tdh2-GFP retained its full length and subcellular localization after evolution (Fig. S2B and S2C), and that there were no mutations in its promoter or coding regions. These data indicate that the increased noise in the evolved line is not caused by mutations in the *TDH2* locus.

If the evolved mutations in the Tdh2-GFP-carrying line occurred within sequences pertaining to general noise regulators, these mutations may affect both the *TDH2*-related pathway and other pathways. We selected four pertinent genes—*TDH3*, *PGK1*, *ADK1*, and *GLY1*—to investigate the effect of the evolved mutations. *TDH3* is a paralog of *TDH2* that encodes an enzyme for glycolysis/gluconeogenesis. *PGK1* encodes another key enzyme involved in glycolysis/gluconeogenesis. Both *TDH3* and *PGK1* are likely co-regulated with *TDH2* since all three respective proteins operate in the same metabolic pathway [28]. *ADK1* encodes an adenylate kinase required for purine metabolism [29], and *GLY1* encodes a threonine aldolase involved in glycine biosynthesis [30], neither of which is related to the *TDH2* pathway. Nonetheless, all four genes exhibited increased protein noise in the evolved line (Fig. 3A), suggesting that the effect of the evolved mutations is not pathway-specific.

**Fig. 3.**
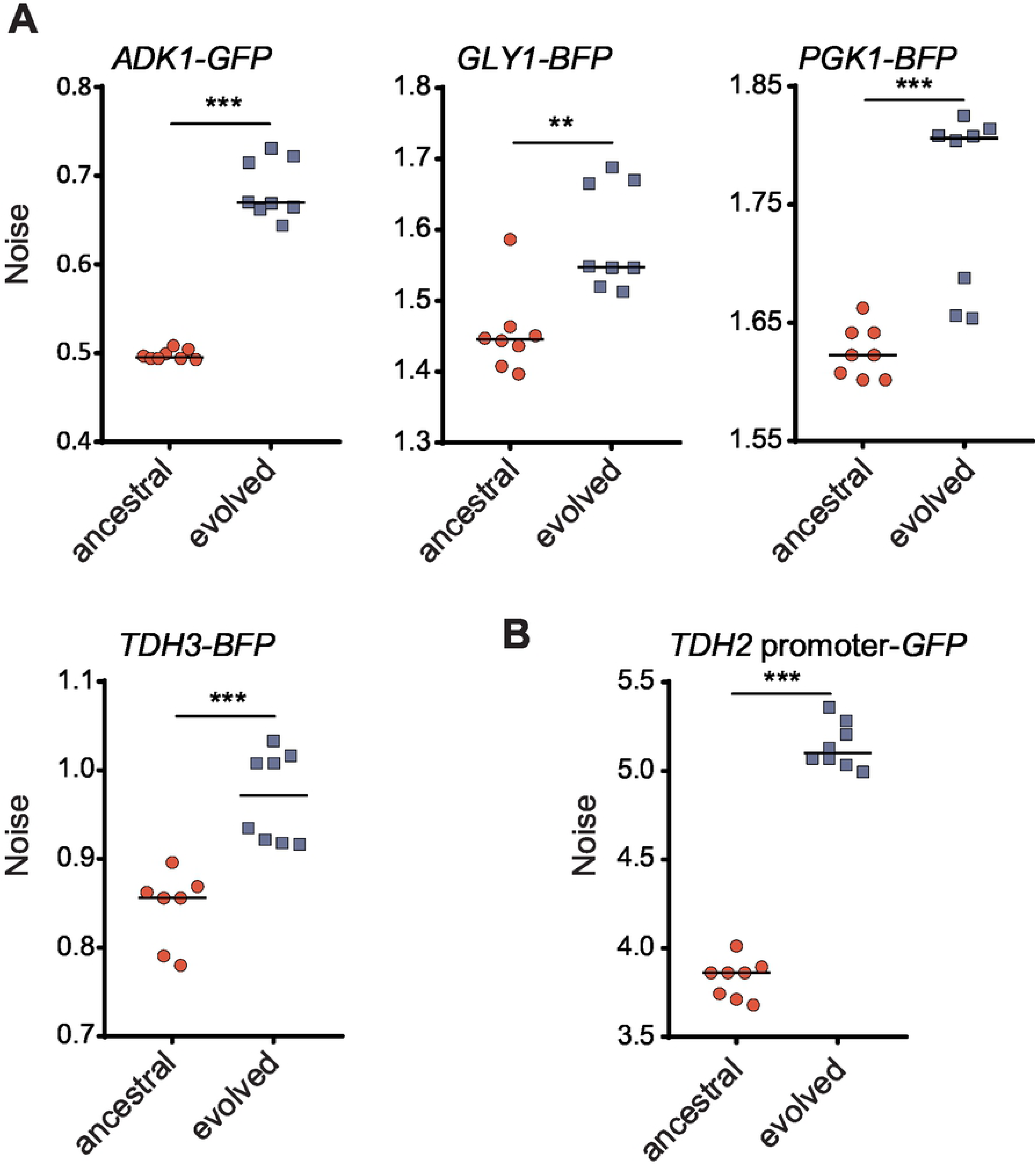
Multiple genes in the evolved cells exhibit increased noise levels. (A) Four reporter genes (*TDH3-BFP*, *PGK1-BFP*, *ADK1-GFP* and *GLY1-BFP*) were engineered in modified ancestral (red circles) and evolved (blue squares) *TDH2-GFP*-carrying cells (see Materials and Methods), and their expression noise was measured (one-sided Wilcoxon rank-sum test, n=7-8; *p* = 4.534 × 10^-4^ for *ADK1*, *p* = 0.0027 for *GLY1*, *p* = 9.441 × 10^-4^ for *PGK1*, *p* = 7.158 × 10^-4^ for *TDH3*). *TDH3* and *PGK1* are involved in *TDH2*-related pathways, whereas *ADK1* and *GLY1* are not. (B) Transcriptional regulation is responsible for increased noise in the evolved cells (one-sided Wilcoxon rank-sum test, n=7-8; *p* = 4.495 × 10^-4^). The *TDH2* promoter was directly fused with the coding sequence of *GFP*. This construct was engineered in modified ancestral and evolved *TDH2-GFP*-carrying cells and its expression noise was measured. The median value of replicates is indicated by horizontal solid lines among groups of data points. \*\**p* < 0.01, \*\*\**p* < 0.001..

We also subjected a *TDH2* promoter-driven GFP construct to the same analysis and observed increased noise in the respective evolved line (Fig. 3B), raising the possibility that increased noise in the evolved cell lines may be attributable, at least in part, to promoter regulation or mRNA turnover.

### An Hmt1 methyltransferase mutant significantly contributes to the evolved increase in noise

To understand the genetic basis of the increased noise we observed in the evolved Tdh2-GFP-carrying line, both ancestral and evolved lines were subjected to whole genome sequencing. A total of 1022 mutations (including 494 non-synonymous, 256 synonymous, and 271 intergenic mutations) were identified in the evolved genome (Table S1). This high number of mutations is most likely due to the mutagen treatment we applied during the cycles of selection.

Next, we used bulk segregant analysis to refine the list of candidate mutations. To do this, we crossed the evolved line to the ancestral line and analyzed their F1 haploid progeny. We measured the noise level of Tdh2-GFP in 360 segregants and established an “evolved-like” pool (comprising 16 segregants) and an “ancestral-like” pool (20 segregants; see Fig. S3 and Materials and Methods). Both of these segregant pools were then subjected to whole genome sequencing. Based on our computational simulation, we applied two criteria to select candidate mutations from these segregant pools: 1) the mutation frequency in the “evolved-like” pool had to be >70%; and 2) the difference in mutation frequencies between the “evolved-like” and “ancestral-like” pools was >38% (see Materials and Methods). Twenty non-synonymous mutations met these two assumptions and were subjected to reconstitution experiments (Table S2).

We first introduced the candidate mutations into the ancestral line using the CRISPR/Cas9 system and then examined the noise level of Tdh2-GFP. Among the tested reconstitution lines, we found that a mutation (G70D) in the methyltransferase-encoding gene, *HMT1*, resulted in a significant increase of noise, close to the level exhibited by the evolved line (Fig. 4A). Similarly, we observed increased noise in the *TDH2* promoter-driven GFP construct (Fig. 4B). When we rescued the mutation in the evolved line by reverting to the wild-type sequence, Tdh2-GFP noise was reduced to a level similar to the ancestral line (Fig. 4A). These data indicate that *hmt1-G70D* is a primary mutation contributing to the evolved increase in noise.

**Fig. 4.**
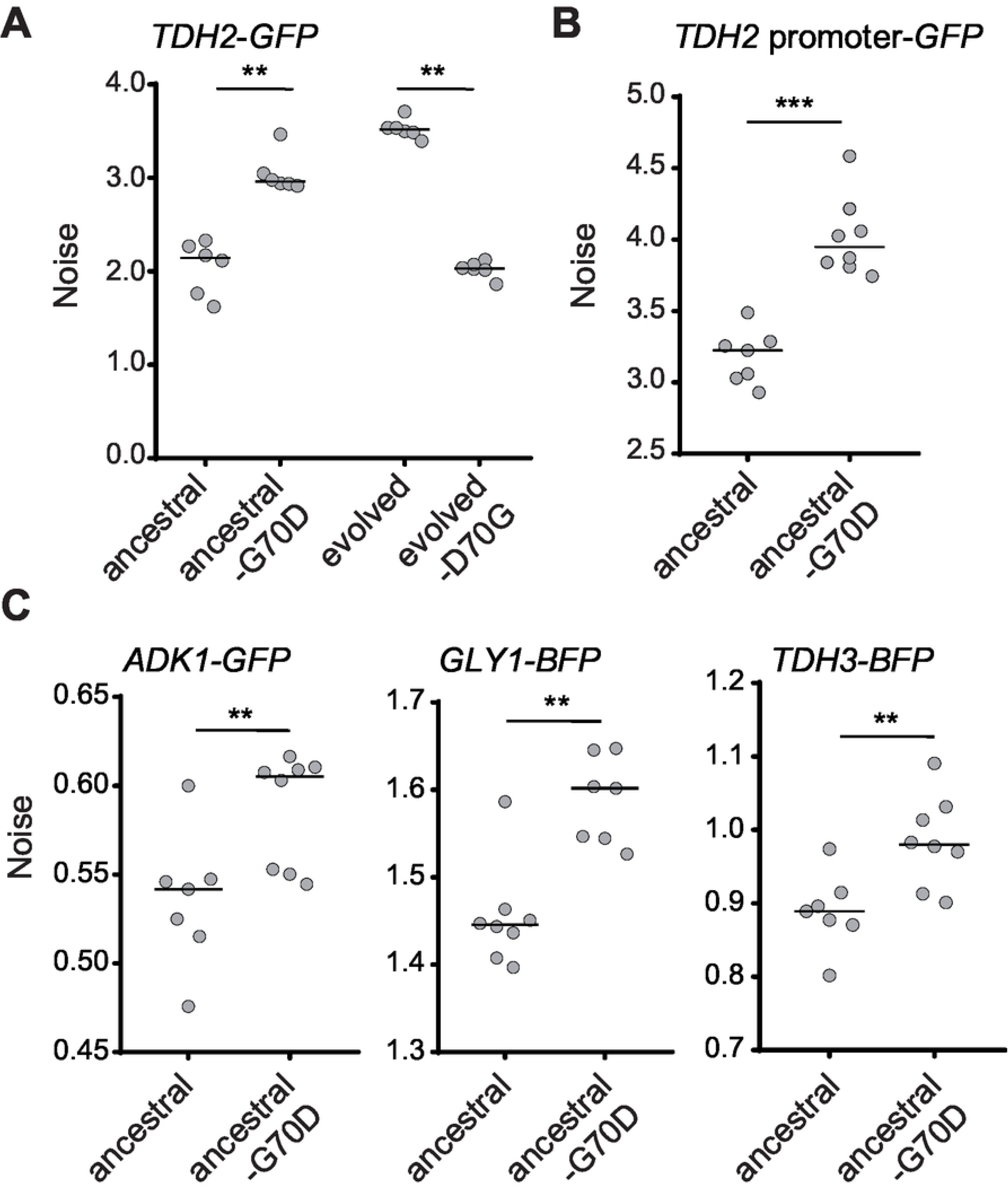
The *hmt1-G70D* mutant recapitulates the increased noise phenotype observed in the evolved line. (A) Reconstituting the *hmt1-G70D* mutation in the ancestral background increases Tdh2-GFP noise, whereas reversing that mutation in the evolved background decreases it (one-sided Wilcoxon rank-sum test, n=6; *p* = 0.0011 for the ancestral background, *p* = 0.0025 for the evolved background). (B) Reconstituted *hmt1-G70D* mutant cells exhibit increased expression noise of the *TDH2* promoter-GFP construct (one-sided Wilcoxon rank-sum test, n=8; *p* = 0.0002). (C) Reconstituted *hmt1-G70D* mutant cells present increased noisy expression of Adk1-GFP, Gly1-BFP, and Tdh3-BFP (one-sided Wilcoxon rank-sum test, n=7-8; *p* = 0.0030 for *ADK1*, *p* = 0.0011 for *GLY1*, *p* = 0.0030 for *TDH3*). All mutants were constructed using the CRISPR/Cas9 system. The median value of replicates is represented by horizontal solid lines among groups of data points. \*\**p* < 0.01, \*\*\**p* < 0.001.

Already in this study we have shown that proteins in both *TDH2*-related and unrelated-pathways exhibited increased noise in the evolved line (Fig. 3A). In the *hmt1-G70D* reconstitution line, we observed similar noise increases in the *TDH3*, *ADK1*, and *GLY1* reporter genes (Fig. 4C), indicating that the increased noise observed in the evolved line is mainly due to the *hmt1-G70D* mutation.

### The noise buffering effect of Hmt1 is mediated through multiple methylation targets

Hmt1 is a methyltransferase that methylates arginine residues in its substrates. The mutated G70D residue is located within a conserved methyltransferase motif (Fig. S4A), and mutations in this motif cause loss of methyltransferase activity in *E. coli* and yeast [31, 32]. The *hmt1-G70D* mutant also exhibited a noise increase similar to levels exhibited by *HMT1*-deletion cells (Fig. S4B). Moreover, Western blot analysis using anti-methylated arginine antibodies confirmed that *hmt1-G70D* mutant cells lack Hmt1 methyltransferase activity (Fig. S4C). These results suggest that the noise regulating function of Hmt1 is mediated through its methyltransferase activity.

In vivo and in vitro studies have identified several Hmt1 substrates [33–35]. Moreover, Hmt1-mediated methylation has been shown to enhance the function of its substrates in multiple pathways, including chromatin remodeling and transcription [36–38], translation and ribosome biogenesis [39–42], and post-transcriptional regulation [43–46]. We selected the representative proteins Npl3, Rps2, Sbp1, and Snf2 from among these pathways and generated corresponding deletion or hypomorphic mutants (in cases where mutant haploids died or had severe growth defects) and investigated noise levels among these mutants.

Of all these tested mutants, only the *rps2* and *snf2* mutants presented significantly increased noise (Fig. 5A), with the other mutants showing either no change or reduced noise (Fig. 5B). Rps2 is a component of the small ribosomal subunit, suggesting involvement of translational regulation in noise control. Snf2 is the catalytic subunit of the SWI/SNF chromatin remodeler and its function depends on two other SWI/SNF components, Snf5 and Snf6 [47, 48]. We further assessed *snf5* and *snf6* mutants and found that they also presented increased noise (Fig. 5A and 5C), confirming the general role played by the SWI/SNF chromatin remodeler in noise regulation.

**Fig. 5.**
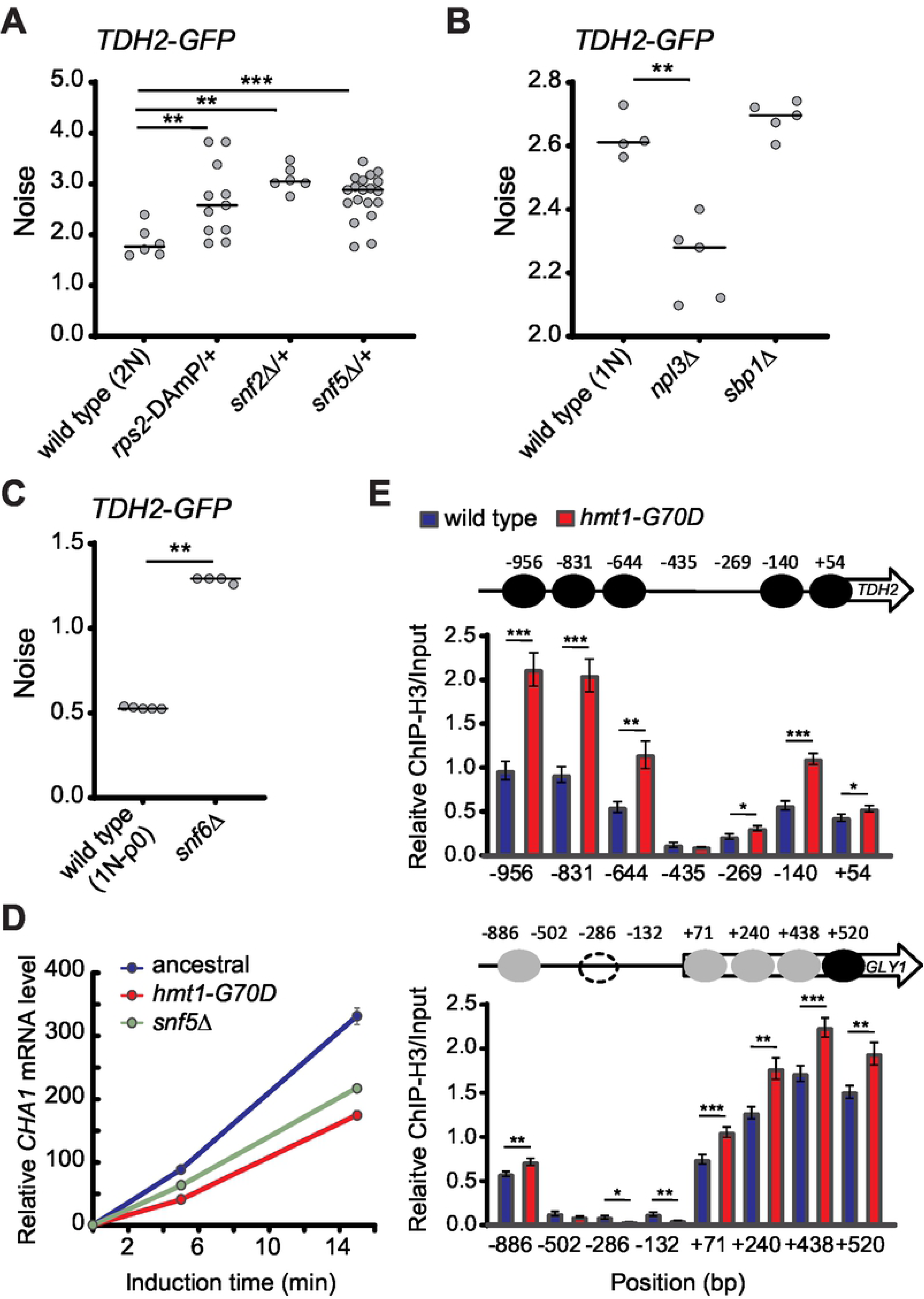
Mutations of SWI/SNF components and the small ribosomal subunit Rps2 result in elevated noise. (A) Attenuating *RPS2*, *SNF2*, or *SNF5* gene expression in the ancestral background results in increased noise (one-sided Wilcoxon rank-sum test, n=6-19; *p* = 0.0039 for *rps2*-DAmP/+, *p* = 0.0025 for *snf2*Δ/+, *p* = 0.0008 for *snf5*Δ/+). Rps2 and Snf2 are methylation substrates of Hmt1. Snf2, Snf5, and Snf6 are essential components of the SWI/SNF chromatin remodeler. Since haploid mutants of *rps2*-DAmP, *snf2*Δ, and *snf5*Δ exhibit severe growth defects and as slow growth has been shown to increase expression noise [86], we constructed heterozygous mutant diploids to measure noise. (B) Deletions of *NPL3* or *SBP1* in haploid cells do not result in increased noise (one-sided Wilcoxon rank-sum test, n=5; *p* = 0.0079 for *npl3*Δ, *p* = 0.206 for *sbp1*Δ). Npl3 and Sbp1 are methylation substrates of Hmt1. (C) Deleting *SNF6* in the ancestral background results in increased noise (one-sided Wilcoxon rank-sum test, n=5; *p* = 0.0087). Haploid *snf6*Δ mutants exhibit defective respiration, so we used a haploid rho-ancestral strain in this experiment and cells were grown in YPD medium. The median value of replicates is represented by horizontal solid lines among groups of data points. (D) The activity of SWI/SNF chromatin remodeling complexes is compromised in the *hmt1-G70D* mutant. It has been shown previously that compromising SWI/SNF complexes results in attenuated transcriptional activation of *CHA1* [49]. Total RNA was isolated from ancestral, *hmt1-G70D,* and *snf5*Δ haploid cells 0, 5, and 15 min after adding 0.1% L-Serine (an inducer of *CHA1* expression). Specific mRNA levels were assessed by Q-PCR. In the figure, *CHA1* mRNA levels were normalized to those of *PYK1*, with this latter acting as an internal control. For all time-points, *CHA1* mRNA levels in ancestral cells (blue) were significantly higher than those in *hmt1-G70D* (red) or *snf5*Δ (green) mutant cells (one-sided Student’s t-test, n = 9; *p* < 0.01). (E) *hmt1-G70D* mutant cells display higher nucleosome occupancy in the promoter and partial coding regions of *TDH2* and *GLY1*. The chromatin status of the *TDH2* promoter was established by chromatin immunoprecipitation against histone 3 coupled with Q-PCR. The numbers in the x-axis indicate the distance (in bp) from the transcription start site. The y-axis represents relative enrichment of histone 3 signals for the amplicons at the indicated regions (one-sided Student’s t-test, n = 10-12). Black, grey, and dashed circles indicate confirmed, fuzzy, and condition-specific nucleosome-occupied regions, respectively. Error bars represent standards errors. \**p* < 0.05, \*\**p* < 0.01, \*\*\**p* < 0.001.

To test whether *hmt1-G70D* mutation affects the activity of SWI/SNF complexes, we performed a functional assay of the SWI/SNF chromatin remodeler. It has been shown previously that transcriptional activation of the *CHA1* gene is attenuated when cells have defective SWI/SNF complexes [49]. We monitored mRNA levels of *CHA1* immediately after shifting cells to a *CHA1*-inducing medium. Expression of *CHA1* was indeed reduced in *hmt1-G70D* mutant cells and in *snf5* mutants (Fig. 5D), indicating that the function of SWI/SNF complexes is compromised in *hmt1-G70D* mutant cells.

We then examined the promoter regions of *TDH2* and *GLY1* (unrelated to *TDH2*) by histone chromatin immunoprecipitation (ChIP) combined with quantitative PCR to understand how Hmt1 influences gene expression. We observed increased nucleosome occupancy in *hmt1-G70D* mutant cells (Fig. 5E), providing evidence for the supposition that noise regulation partially operates via SWI/SNF-mediated nucleosome remodeling [18].

### Hmt1 functions as a mediator in responses to environmental stresses

Genome-wide studies have indicated that *HMT1* expression frequently fluctuates under different growth conditions [50, 51], suggesting that cells may employ an environmental sensor to enhance phenotypic heterogeneity upon encountering stress. We measured the mRNA levels of *HMT1* under various stress conditions—including heat, oxidative stress, high osmolarity, and glucose starvation—and found that they were significantly reduced under all of these conditions (Fig. 6A). Consistent with the concept of an environmental sensor, the noise level of Tdh2-GFP was also increased under stress conditions (Fig. 6B and S5). Together, these data indicate that Hmt1 can serve as a mediator to control levels of phenotypic heterogeneity in response to environmental stimuli.

**Fig. 6.**
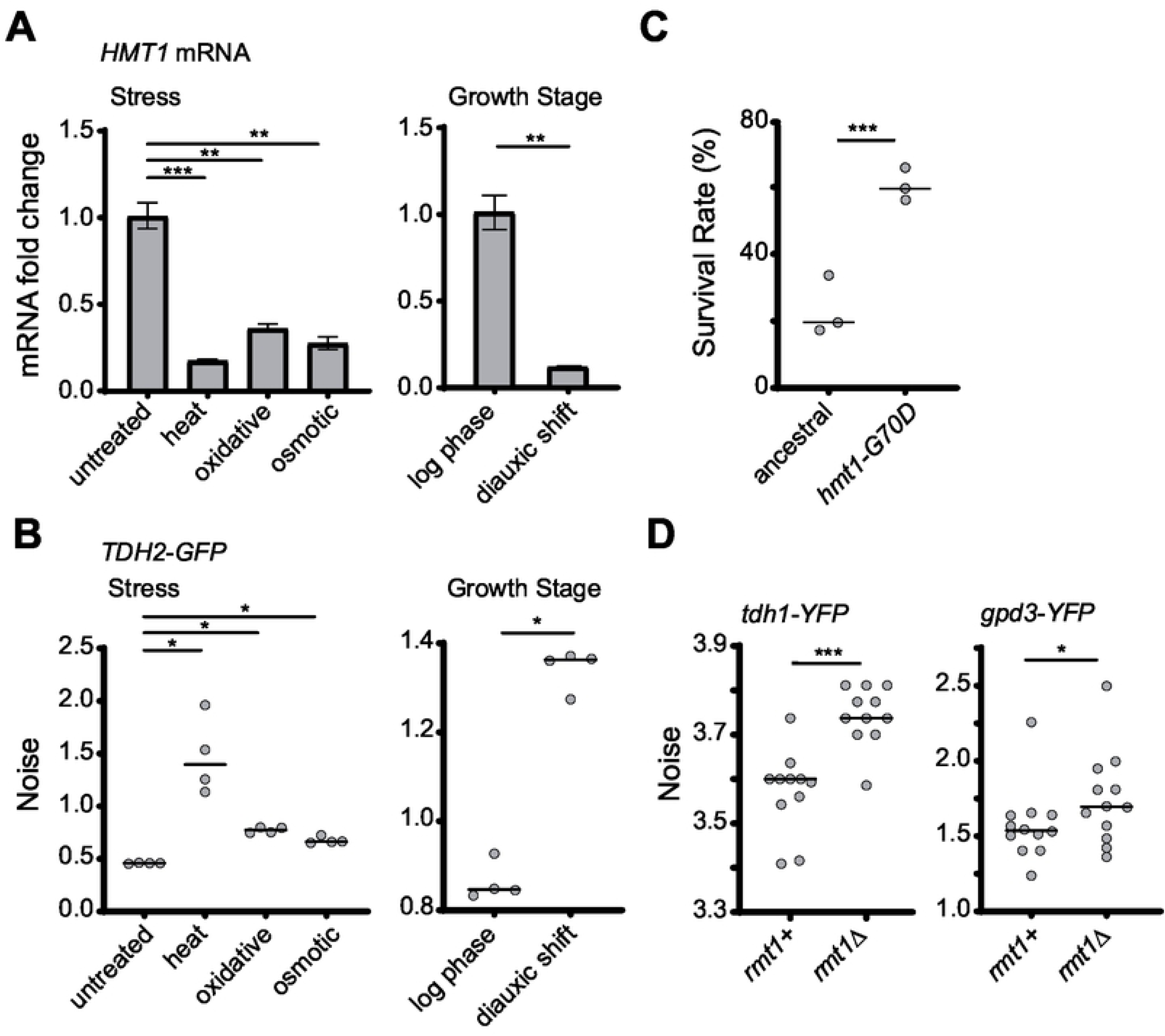
Hmt1-mediated noise suppression can be released under non-optimal growth conditions as a conserved cell survival strategy. (A) *HMT1* transcripts are down-regulated under non-optimal growth conditions (one-sided paired Student’s t-test, n = 4; *p* = 0.0008 for heat stress, *p* = 0.003 for oxidative stress, *p* = 0.002 for osmotic stress, *p* = 0.007 for diauxic shift). The level of mRNA was measured using Q-PCR, and *DET1* mRNA was used as the internal control. Error bars indicate standard errors. (B) Non-optimal growth conditions result in increased Tdh2-GFP noise (one-sided Wilcoxon rank-sum test, n = 4; *p* = 0.015 for heat, oxidative and osmotic stress, *p* = 0.014 for diauxic shift). Noise was measured 2.5 h after cells had been shifted to the indicated conditions or was measured at different growth stages. (C) *hmt1-G70D* mutant populations survive better than the ancestral line in medium containing H_2_O_2_ (Fisher’s exact test; *p* < 2.2 x 10^-16^). Stationary-phase cells were spread on plates with 5.3 mM H_2_O_2_ and survival rates were determined by counting colony-forming units after 5 days of growth. (D) Mutation of the *HMT1* ortholog in the fission yeast *S. pombe* also results in increased expression noise. *rmt1* is the ortholog of *HMT1*, whereas *tdh1* and *gpd3* are *TDH2* orthologs. Deletion of *rmt1* increased protein noise of both tdh1-YFP and gpd3-YFP (one-sided Wilcoxon rank-sum test, n = 10-12; *p* = 0.0005 for *tdh1-YFP*, *p* = 0.037 for *gpd3-YFP*). The median value of replicates is indicated with horizontal solid lines among groups of data points. \**p* < 0.05, \*\**p* < 0.01, \*\*\**p* < 0.001.

Our data show that bet-hedging within a population could be represented by the noisy expression of Tdh2-GFP (Fig. 2). If increased noise can help populations survive stressful environments, we anticipated that cells with low Hmt1 activity under stress would present an enhanced survival rate relative to those with normal Hmt1 activity under stress. Indeed, when we treated wild-type and *hmt1-G70D* mutant cells with H_2_O_2_, the mutant population exhibited better viability (Fig. 6C).

### Hmt1-mediated noise buffering is conserved in *Schizosaccharomyces pombe*

Our results suggest that the methyltransferase Hmt1 may function as a general noise regulator that constrains physiological noise in normal conditions but facilitates heterogeneity under stress. This buffering mechanism is likely to increase long-term population survival. Methyltransferase is a conserved enzyme that exists in all eukaryotic kingdoms. We examined whether its buffering function is also conserved in a phylogenetically distinct species, *S. pombe*, which diverged from the common ancestor of *S. cerevisiae* at least 400 million years ago [52]. We generated YFP fusion protein constructs of *tdh1* and *gpd3* (the *S. pombe* orthologs of *TDH2*) and examined their noise levels in the wild-type and *rmt1* (the *S. pombe* ortholog of *HMT1*) deletion backgrounds. Similar to our findings for *S. cerevisiae* evolved lines, noise levels of Tdh1-YFP and Gpd3-YFP were significantly increased in *rmt1* mutant cells (Fig. 6D). Accordingly, the Hmt1-mediated noise buffering system probably represents an important survival strategy that is conserved across diverse microorganisms.

## Discussion

Microorganisms constantly face changing environments. Despite being equipped with complex stress-adaption systems, unpredicted acute stress remains a challenge for cells. Cell populations harboring heterogeneous physiological states enhance their likelihood of surviving environmental fluctuations [53, 54]. Many examples of bet-hedging have been reported previously, indicating that this is a common survival strategy among microbes [5, 55]. However, it is not known if cells exhibit another layer of regulation that allows them to adjust their levels of noise. Our current study provides evidence that Hmt1 can function as a core regulator to constrain or facilitate phenotypic heterogeneity in response to environmental stimuli.

The evolutionary advantage of an environment-sensing noise regulator is readily conceivable. Although stochastic noise is inevitable among individuals within a population, excessive deviation from “normal” levels may result in a significant reduction of fitness under normal conditions. By regulating multiple pathways, Hmt1 allows cells to constrain noise levels. However, when a population is exposed to mild stresses, Hmt1 expression is immediately down-regulated and the noise buffering system is curtailed. The resulting enhanced heterogeneity means that the population increases its likelihood of survival, especially if the stress is prolonged or escalates (Fig. 7).

**Fig. 7.**
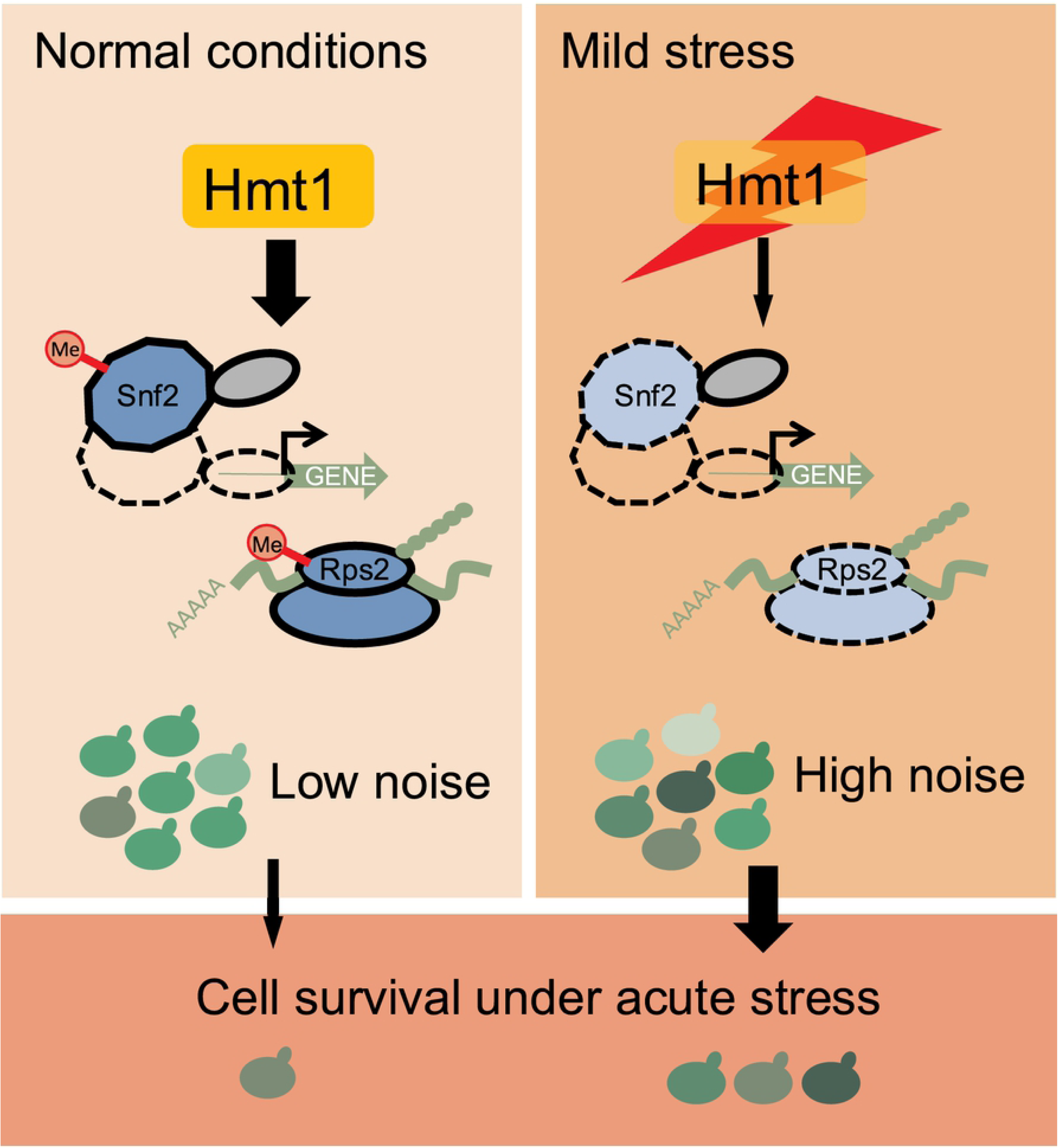
A model showing how Hmt1 modulates cell-to-cell heterogeneity in response to environmental stress. Hmt1 methylates and enhances the function of the SWI/SNF chromatin remodeler and small ribosomal subunits to reduce stochastic noise in gene expression. Under normal conditions, cells maintain a high level of Hmt1 and exhibit homogeneous gene expression in most cells. However, when the population encounters environmental stress, *HMT1* expression is down-regulated, inhibiting the functions of Hmt1 targets. Accordingly, expression of Hmt1 gene targets becomes noisier so individual cells exhibit heterogeneous cell physiologies. The likelihood of population survival is enhanced due to this heterogeneity.

Hmt1 is a type-I arginine methyltransferase that catalyzes mono-methylation and asymmetric di-methylation in budding yeast. Hmt1-mediated methylation has been shown to influence various cellular pathways including transcription, post-transcription and translation [36-38, 40, 42-46]. Moreover, Hmt1 is a highly interactive protein, ranking among the top 1.5% of the entire budding yeast proteome in terms of the interaction number [56], suggesting it has a central role in coordinating a complex network. Previous studies have shown that another network hub, Hsp90, can regulate general cellular noise [17, 57]. Consistent with predictions from network analyses and results from a genome-wide screen, hub genes have a strong impact on buffering non-genetic and genetic variation [58–60]. More interestingly, both Hsp90 and Hmt1 can work as environmental sensors to adjust noise levels in response to environmental cues, acting as direct links between noise regulation and adaptive benefits.

Our results show that at least two downstream pathways, i.e., chromatin remodeling and the translational machinery, are involved in Hmt1-mediated noise buffering. The effect of the SWI/SNF chromatin-remodeling complex on gene noise has been reported previously [11]. Here, we have identified Hmt1 as an upstream coordinator of noise regulation and establish the novel role of protein methylation in this process. The detailed mechanism underlying how the ribosomal component Rps2 regulates noise is less clear. Rps2 plays a crucial role in controlling translational accuracy and its efficiency can be further modulated by post-translational modifications [61, 62]. It is possible that Rps2-mediated noise regulation occurs at the protein translation level. More experiments are required to address this issue.

The regulatory functions of Hmt1 are generally well conserved in human cells, but they are executed by six orthologs [33, 63]. Human orthologs also exist for many Hmt1 substrates, which interact respectively with different human type I enzymes [34, 64–66]. For example, the human Hmt1 ortholog PRMT4 interacts with the Snf2 ortholog BRG1 to facilitate the ATPase activity of the entire human SWI/SNF complex [66]. It is likely that the interactions between Hmt1 and its substrates represent an ancient regulatory network. In support of this supposition, we observed increased noise in *S. pombe* and *S. cerevisiae hmt1* mutants, suggesting that Hmt1-mediated buffering evolved hundreds of millions of years ago before the ancestors of these two species diverged. Hmt1-mediated noise buffering may also help populations of unicellular *S. pombe* to survive in stressful environments. However, with regard to multicellular organisms, methylation-regulated noise buffering may influence their developmental plasticity, and this interesting topic awaits further study.

## Materials and Methods

### Yeast strains and genetic procedures

All *Saccharomyces cerevisiae* strains used in this study were derived from the W303 strain (*leu2-3, 112 trp1-1 can1-100 ura3-1 ade2-1 his3-11, 15*), and the *Schizosaccharomyces pombe* strains were derived from the 972 h^-^ strain. Unless otherwise indicated, gene deletion or insertion was based on homologous recombination. Yeast cells were transformed using the Lithium acetate method [67] or electroporation under 1800 V/ 200 Ω/ 25 μF (BTX Gemini SC^2^, Fisher Scientific, Pittsburgh, PA) [68].

All fluorescent fusion proteins were constructed by chromosomal in-frame insertion of the fluorescent protein tag at the 3’ end of the coding region of target genes. The GFP tags were directly amplified from the genomic DNA of corresponding strains in the yeast GFP collection [69]. The BFP tag was amplified from the plasmid pFA6a-link-yomTagBFP2-Kan [70]. Prior to constructing *ADK1*-*GFP* and *TDH2* promoter-driven *GFP* in ancestral and evolved *TDH2-GFP*-carrying lines, the *GFP* of *TDH2*-*GFP* in these strains was removed and replaced with a stop codon and the *TDH2* 3’ untranscribed region (UTR). For *TDH2* promoter-driven *GFP*, 1000 base pairs (bp) upstream of the *TDH2* coding region was fused with *GFP*, and the fused fragment was inserted between positions 204886 and 204887 of chromosome I [71]. For YFP-tagging in *S. pombe*, the coding region of yVenus [72] without the start codon was fused with hphMX6 by two-fragment PCR. To delete *S. cerevisiae* genes, KanMX4-containing DNA fragments for homologous recombination were amplified from the genomic DNA of corresponding strains in the yeast deletion collection [73]. To construct a hypomorphic mutant of *RPS2*, we directly amplified the *rps2*-DAmP allele from the yeast DAmP diploid collection [74] and used it to replace the native gene. To delete *rmt1* in *S. pombe*, a KanMX6-containing DNA fragment with homologous flanking regions was generated and used to replace the whole gene.

All SNP mutant reconstitution strains were constructed using the CRISPR/Cas9 system [75, 76]. Briefly, host cells were transformed with the Cas9 expression plasmid and then the transformant was introduced with a DNA fragment encoding gRNA, a linearized vector, and mutation-containing donor DNA fragments. After transformation, single colonies were streaked out on YPD (1% <u>Y</u>east extract, 2% <u>P</u>eptone and 2% <u>D</u>extrose) plates to purify the CRISPR transformants. Genomic DNA of the resulting single colonies was isolated and examined initially by allele-specific PCR [77] and then by Sanger sequencing. The plasmids for Cas9 and gRNA expression are listed in Table S3.

### Experimental evolution

For the evolution experiment, eight genes (*ADK1*, *APA1*, *PCM1*, *RPL4B*, *SAM4*, *TDH2*, *TPD3*, and *TYS1*) were selected and fused with GFP to generate evolving strains each carrying an individual reporter gene. These reporter genes were chosen based on the following criteria: 1) each gene belongs to a distinct cellular pathway; 2) the expression level of GFP fusion proteins is constant during the cell cycle and is sufficiently high to be detectable by flow cytometry; and 3) the reporter proteins are evenly localized in the nucleus or cytosol to avoid misinterpretation due to dynamic organelles.

Due to the complexity of our selection regime, only one evolving line for each reporter gene was established. In each selection cycle, 1 × 10^6^ cells were treated with 2.8% EMS (see the “EMS mutagenesis” section below for details) to increase the genetic diversity of the cell population. This mutagenic treatment is crucial, as we initially ran a pilot experiment using a similar selection regime but lacking the mutagen treatment and did not observe any obvious increase in noise after 70 cycles of selection. However, our EMS treatments also significantly increased the number of mitochondria-defective cells, which are known to increase population heterogeneity. To constrain the population of mitochondria-defective cells, we grew cells in non-fermentable YPG (1% <u>Y</u>east extract, 2% <u>P</u>eptone and 2% <u>G</u>lycerol) medium at 28 °C after EMS treatments. After 12 h of growth in YPG, cells were sorted to select 5000 cells from the top (or bottom) 5% of the total population in terms of GFP intensity (see the “Fluorescence-activated cell sorting” section below for details). These cells were grown in 3 ml YPG for ∼36 h to reach OD_600_ = 1 before proceeding to the next cycle of selection. We used alternating selection between the top 5% and bottom 5% of total populations throughout the evolution experiment. The effective population size was estimated to be 1.33 × 10^5^ cells using the formula 2/*N_e_*= 1/(*N_01_* × *g*) + 1/(*N_02_* × *g*), in which *N_0_* is the initial population size and *g* is the number of generations during each growth period [78].

After 35 cycles of selection, five individual clones were isolated from each evolved population and their GFP noise levels were measured. The clone with the greatest increase in noise without exhibiting a decrease in the mean intensity of GFP or increased variation in cell size was selected for further genetic analysis.

### Ethyl methanesulfonate (EMS) mutagenesis

Mutagenesis was performed according to a previously published protocol by which mutation rates can be increased without inducing considerable cell death [79]. Briefly, we washed 1 × 10^6^ cells with sterile water once, with 100 μl of phosphate buffer (0.1M Na_2_HPO_4_, pH=7.0) once, and then resuspended them in 90 μl of phosphate buffer. We added 90 μl of EMS-containing phosphate buffer to the cell solution, resulting in a final concentration of 2.8% EMS. The solution was maintained under constant shaking at room temperature for 30 min, before stopping the reaction by adding 50 μl of 25% sodium thiosulfate. After washing with sterile water, the cells were transferred to 10 ml YPG. The survival rate of wild-type cells after EMS treatment ranged from 60% to 80%.

### Fluorescence-activated cell sorting and flow cytometry

Cells were suspended in filtered phosphate-buffered saline (PBS) (137 mM NaCl, 2.7 mM KCl, 10 mM Na_2_HPO_4_, and 1.8 mM KH_2_PO_4_) and sampled using a BD FACSJazz™ machine (BD Bioscience, Franklin Lakes, NJ) at an event rate of ≤ 3000 cells/sec. GFP readouts were acquired with bandpass filters of 513/17 nm using laser excitation at 488 nm. For YFP, we used laser excitation at 488 nm with a 542/27 nm filter and for BFP we used a 450/50 nm filter and laser excitation at 405 nm.

To eliminate interference from small particles in the fluidic system, we excluded particles with forward scatter (FSC) signals <2570 (as a trigger threshold). We established three hierarchical gates. First, to avoid cell aggregates, we collected cells that had relatively constant signals of trigger pulse width along FSC signals. Second, to eliminate possible cell aggregates, we collected cells that had relatively constant signals of FSC-W along FSC signals. Third, to avoid cell debris and abnormally large cells, we collected cells having FSC and side scatter (SSC) signals both ranking within 5%-95% of the population.

For alternating selection in experimental evolution, we applied the trigger threshold and the first and third gates. Based on the distribution of GFP levels, we defined the highest or lowest 5% of the total gated population and collected 5000 cells. The collected cells were then grown in 3 ml YPG and used for the next run of selection. For fitness assays, we applied the trigger threshold and all three gates. Based on the distribution of GFP levels, we defined the highest and lowest 5% of the total gated population and collected individual cells. A 1.0 single-drop sorting mode was employed. The viability of the collected cells was then measured under different conditions.

### Noise and signal distribution measurement

We measured the noise of fluorescent protein signals by the Fano factor (a ratio of variance to the mean, %) using more than 5000 early log-phase cells. This measurement is characterized by having less interference from the mean and is more sensitive to variation-driven increased noise [11, 80]. We used the same trigger threshold and the first two hierarchical gates as employed for cell sorting in our noise measurement. For the third gate, we used contour plots of FSC and SSC signals to collect cells constituting 60% of the second-gated population to ensure more homogenous cell size and cell physiology. Fluorescence readouts of the entire population were log-transformed and used to calculate the noise.

### Fitness assays under different growth conditions

To measure H_2_O_2_ resistance, Tdh2-GFP-carrying cells were grown in YPD for 5 days to ensure that the cells had entered stationary phase. More than 100 cells from the top 30% or bottom 30% of the gated population in terms of their Tdh2-GFP intensity were sorted and spotted onto plates with or without 4.4 mM H_2_O_2_. The survival rates were measured after 5 days of incubation at 28 °C.

For heat resistance assays, cells from the top 10% or bottom 10% in Tdh2-GFP intensity were sorted into microcentrifuge tubes. The collected cells were divided into two parts. One part was immediately placed in a PCR machine to perform heat ramping from 30 °C to 56 °C for 20 min as a heat stress [81]. The other part was placed in fresh YPD media for 2.5 h and then subjected to the same heat treatment. Cell survival rates were measured as colony-forming units after 3 days of incubation at 28 °C.

As a rebudding assay, more than 100 unsorted stationary-phase cells were placed on agarose pads and monitored by time-lapse microscopy at 15 min intervals for at least 12 h [82]. Only unbudded cells at the beginning of the recording period were monitored.

### Western blotting

Log-phase cells were lysed using NaOH lysis [83]. Proteins in the lysates were separated with SDS-PAGE and transferred to a PVDF membrane. Mouse anti-GFP antibody (1:4000) (#sc-9996, Santa Cruz Biotechnology, Dallas, TX) was used to detect Tdh2-GFP. Rabbit anti-G6PDH antibody (1:4000) (#A9521, Sigma-Aldrich, St. Louis, MO) was used to detect G6PDH, which served as an internal control. Mouse anti-methylated arginine antibody (mab0002-P, Covalab, Villeurbanne, France) was used to detect proteins with methylated arginine. MultiMab rabbit monoclonal mix antibodies against an asymmetric di-methyl arginine motif (#13522, Cell Signaling Technology, Danvers, MA) were used to detect proteins with asymmetric di-methylated arginine.

### Microscopy

Microscopy was conducted using a 60x objective lens and an ImageXpress Micro XL system (Molecular Device, Sunnyvale, CA).

### F1 segregant analysis

The evolved line was mated with the ancestral line, and the resultant diploid cells were induced to sporulate. Sporulated culture was harvested into a microcentrifuge tube and treated with 0.5 mg/ml Zymolyase-100T (Nacalai Tesque Inc., Kyoto, Japan) in 1M sorbitol at 28 °C for 2 h to remove the ascal wall. Cells were then treated with 2% SDS at 28 °C for 10 min to kill unsporulated diploid cells, before washing with sterile water and vortexing vigorously to attach individual spores to the tube wall. We then added 0.01% Triton X-100 solution to the tube, before vigorous sonication to detach spores from the tube wall and to separate spore clusters. The suspension was diluted and spread on YPD plates to isolate F1 haploid segregants.

We conducted a total of three runs of noise measurement to identify “evolved-like” and “ancestral-like” F1 segregants. The progenies ranking within the top 20% or bottom 20% of the Tdh2-GFP noise level without changing the mean intensity (i.e., within three standard deviations of the control) were selected for the next run of noise measurement. We started with 360 F1 progeny of confirmed ploidy in the first round, and obtained 16 “evolved-like” segregants and 20 “ancestral-like” segregants after finishing the third run. These cells were subjected to whole genome sequencing analysis.

### Whole genome sequencing analysis

Yeast cells were resuspended in 200 μl of lysis buffer (2% Triton X-100, 1% SDS, 100 mM NaCl, 10 mM Tris-HCl pH=8.0, and 1 mM Na_2_EDTA). Glass beads (0.5 mm in diameter) that amounted to the volume of the cell pellet were added, followed by the addition of 200 μl of PCIA (Phenol:chloroform:isoamyl alcohol=25:24:1, pH=8.0). The mixture was vortexed for 5 min. We then added 200 μl of TE buffer (10 mM Tris·Cl; 1 mM Na_2_EDTA pH=8.0) and the mixture was centrifuged for 5 min at maximum speed. Only 350 μl of the solution in the aqueous layer was extracted and subjected to EtOH precipitation. The pellet was air-dried and then dissolved in 400 μl of TE buffer containing 75 μg/ml RNase A. The mixture was then incubated at 37 °C for 5 min. We then added 200 μl of PCIA and we inverted the tubes to mix. EtOH precipitation was then performed at room temperature under constant shaking for 2 h. The pellet was air-dried and then dissolved in 100 μl of sterile double-distilled water. The ratio of OD_260_ to OD_280_ of the purified DNA ranged from 1.8 to 2.1. Concentrations of genomic DNA were determined using a Qubit™ dsDNA BR assay kit (ThermoFisher Scientific).

Equal amounts of genomic DNA from individual “evolved-like” or “ancestral-like” F1 segregants were pooled. These two DNA pools were sequenced using an Illumina NextSeq500 system (Illumina, San Diego, CA) with 150 bp paired-end reads from 350-bp libraries. At least 57X coverage was achieved for sequencing ancestral and evolved clones, and we obtained 150-200X coverage for the pooled segregants. Sequence results were analyzed using CLC Genomics Workbench 9.1 with default settings for read import, read trimming, read alignment, duplicate-read removal, local realignment, single nucleotide polymorphism (SNP) calling (with the slight modification of using a 10% frequency cut-off in the “ancestral-like” pool to retrieve as many mutations as possible since the default is 35%), and SNP annotation. We used the reference genome of S288C (version R64-2-1) from The *Saccharomyces* Genome Database for SNP calling because it is the best annotated.

### Identifying candidate causal SNPs responsible for increased noise

By using the CLC Genomics Workbench 9.1 function “filter by control read” with a criterion of less than 2 control reads, we could eliminate SNPs existing in both ancestral and evolved lines (Table S1). The resulting list of evolved SNPs was then cross-referenced with the list of SNPs from the “evolved-like” pool. We anticipated that candidate causal SNPs responsible for increased noise would be enriched in the “evolved-like” but not in the “ancestral-like” pools. By simulating sequencing results (using Python 2.7) from 150x coverage, we determined the frequencies of SNPs that were randomly segregated into two pools of 10 or 20 individuals to represent SNP enrichment thresholds for increased noise [84]. We identified SNPs with a frequency >70% in the “evolved-like” pool and with a frequency <32% in the “ancestral-like” pool at a 90% confidence interval as being correlated with increased noise (Table S2). The thresholds are represented by 90% confidence intervals for simulations with 10 individuals and at 95% for those with 20 individuals.

### Histone chromatin immunoprecipitation (ChIP) combined with Quantitative PCR (Q-PCR)

Log-phase cells (∼30 OD_600_) were harvested and crosslinked by 1% formaldehyde at 30 °C for 30 min. After quenching with 125 mM glycine, the cell suspension was lysed by three cycles of 5 min beating and 1 min cooling at 4 °C of 0.5 mm glass beads in FA buffer (50 mM HEPES-KOH pH=7.5, 140 mM NaCl, 1% Triton X-100, 0.1% SDS, and 0.1% Deoxycholate Na salt) supplemented with 1x protease inhibitor cocktail (Protease inhibitor cocktail Set IV in DMSO; Merck, 539136) and 1 mM PMSF. We made a hole in the bottom of the lysate-containing tube to allow the lysate to flow through into a collection tube upon centrifugation at 500 x *g* for 3 min at 4 °C. The chromatin fraction was pelleted down by centrifugation at 12000 x *g* for 15 min at 4 °C and washed twice with FA buffer. We transferred 2 ml of the chromatin suspension into a 15-ml centrifugation tube (BIOFIL), avoiding bubbles. We sheared the chromatin using a Bioruptor (Diagenode, Denville, NJ), with 5 cycles of 15 min (30 sec On and 30 sec Off per min) at “High intensity” mode at 4 °C. Ice-cold water was resupplied in the Bioruptor tank after each cycle to maintain the low temperature. The sheared chromatin was cleared by centrifugation at 12000 x *g* for 15 min at 4 °C. One-fifteenth of the cleared chromatin was used as input and stored at −80 °C. The remaining cleared chromatin was transferred into a tube containing pre-incubated Dynabeads^®^ Protein A (#10002D, Invitrogen, Waltham, MA) with histone 3 antibody (ab#1791, Abcam), followed by end-over-end mixing overnight at 4 °C. The beads were anchored with DynaMag™-2 Magnet (ThermoFisher Scientific, Waltham, MA) to remove unbound molecules and were then sequentially washed with 1 ml of FA buffer, 500 mM NaCl-containing FA buffer, DOC buffer (10 mM Tris-HCl, 1 mM Na_2_EDTA pH=8.0, 250 mM LiCl, 0.5% NP-40, 0.5% Deoxycholate Na salt), and TE buffer (10 mM Tris·Cl; 1 mM Na_2_EDTA pH=8.0). The bound molecules were eluted by adding TES buffer (10 mM Tris·Cl; 1 mM Na_2_EDTA pH=8.0, and 1% SDS) and incubating at 65 °C for 20 min, followed by a second elution with TE buffer (65 °C for 10 min). The two eluents were combined. The eluent and thawed input sample were incubated in 0.125 μg/ml RNase A at 37 °C for 30 min, followed by Proteinase K treatment (2 mg/ml Proteinase K at 42 °C for 1 h). The samples were de-crosslinked at 65 °C overnight. DNA was purified using a QIAquick DNA Purification Kit (Qiagen, Hilden, Germany) according to the manufacturer’s instructions, with the modification of a two-cycle wash step using Qiagen PE buffer. We subjected 0.5 ng of DNA to Q-PCR with primers annealing to nucleosome-occupied and -depleted regions of the promoter and coding regions for *TDH2* and *GLY1* (Table S3) [85].

### Q-PCR of mRNA under different growth conditions

For stress conditions, log-phase cells grown in YPD media were divided into four parts. One part was continuously grown in YPD as an untreated sample and the remaining three parts were treated with 0.375 M KCl for 20 min or 0.4 mM H_2_O_2_ for 20 min or subjected to 42 °C for 30 min. For different growth states, total RNA was extracted from the same batch of cultures when cells were in log phase and entering diauxic shift. To investigate the activity of SWI/SNF chromatin remodeling complexes, cells were first grown to log phase in CSM-Serine (Serine-depleted <u>C</u>omplete <u>S</u>ynthetic <u>M</u>ixture). Then L-Serine was added to a concentration of 0.1% to induce *CHA1* expression, before harvesting cultures 0, 5, and 15 min after induction, with the addition of 0.05% NaN_3_ to stop transcription [49].

RNA from 5-10 OD_600_ of cells was extracted and quantified according to a previous report [68] with modifications. Cells were harvested at 4 °C. RNA quality was examined using an Agilent RNA 6000 Nano kit with an Agilent 2100 Bioanalyzer (Agilent Technologies). Two μg of RNA was reverse-transcribed with 0.5 μg oligo(dT)_18_ primer using an Applied Biosystem™ High-Capacity cDNA Reverse Transcription kit (ThermoFisher Scientific). One μl of the reaction products was then used for Q-PCR in an Applied Biosystem™ 7500 Fast Real-Time PCR system (ThermoFisher Scientific) with gene-specific primers (Table S3).

### Statistical analyses

All statistics were performed using R language (http://www.r-project.org/).

## Acknowledgements

We thank members of the Leu lab for helpful discussion and comments on the manuscript. We thank Shao-Win Wang for the *S. pombe* strains. We also thank John O’Brien for manuscript editing, and Po-Hsiang Hung and the IMB Bioinformatics, Genomics and Imaging cores for technical assistance.

## Supplemental Figure Legends

**Fig. S1. Sorted subpopulations of cells exhibiting low or high Tdh2-GFP are not genetically distinct.** Cells were sorted according to their Tdh2-GFP levels into low (red circle) and high (blue circle) subpopulations, each of which constituted 10% of the whole population (grey). These cells were grown in YPD for the indicated periods of time and analyzed by flow cytometry. The distributions of the low (red) and high (blue) Tdh2-GFP subpopulations are superimposed with that of the parental population (grey outline). The vertical dashed lines indicate the mean signal intensity of the parental population.

**Fig. S2. Experimental evolution alters the Tdh2-GFP signal distribution without affecting protein integrity or subcellular localization.** (A) Increased noise in the evolved Tdh2-GFP-carrying line is due to a flattened distribution rather than a bimodal one. (B) Full-length Tdh2-GFP is retained after evolution. Western blots were hybridized using mouse anti–GFP antibody (1:4000) (the upper blot) and by rabbit anti–G6PDH antibody (1:4000) (the lower blot). (C) Protein localization of Tdh2-GFP is not altered by our experimental evolution approach. Cells were imaged using a 60x objective under the FITC channel (the upper panel) or bright field (the lower panel). The scale bar is 5 μm.

**Fig. S3. Tdh2-GFP signal in F1 progeny selected for whole genome sequencing.** For bulk segregant analysis, 360 F1 progeny were derived by backcrossing the evolved clone to the ancestral clone. The noise and mean of Tdh2-GFP signal in individual progeny were analyzed, and a total of three runs of noise measurement were conducted to identify “evolved-like” and “ancestral-like” F1 segregants. Data from the final run of analysis are shown here. gDNA of the “evolved-like” and “ancestral-like” F1 progeny was extracted and respectively pooled for whole genome sequencing.

**Fig. S4. The *hmt1-G70D* mutation phenocopies the loss-of-function mutation.** (A) The identified G70D mutation of Hmt1 is located in a highly conserved methyltransferase motif. An alignment of the primary sequences of the conserved motif is shown for various methyltransferases from budding yeast, fission yeast, human and bacteria. Residues shared with *S. cerevisiae* Hmt1 are labeled in yellow, and the mutated glycine residue observed in the evolved clone is indicated by an arrowhead. (B) Both *hmt1-G70D* and Hmt1 deletion mutants exhibit a similar level of increased Tdh2-GFP noise (one-sided Wilcoxon rank-sum test, n=5-10; *p* = 0.0013 for *hmt1-G70D*, *p* = 0.0013 for *hmt1*Δ). (C) The G70D mutation of Hmt1 results in defective methylation. The patterns of asymmetric di-methylation (left) and pan-methylation (right) of arginine in whole cell lysates of *hmt1-G70D* mutant cells did not differ from those of deletion mutants (*hmt1*Δ), but differed from those of wild type cells.

**Fig. S5. Tdh2-GFP noise significantly increases shortly after stress treatments.** Tdh2-GFP noise increases under non-optimal growth conditions (one-sided Wilcoxon rank-sum test, n=4; *p* = 0.015 for heat stress, *p* = 0.015 for oxidative stress). Noise was measured after cells were treated with the indicated stress for 20-30 min. The difference between untreated and treated cells became more obvious after 2 h (Fig. 6B), which probably reflects the time it takes for cells to alter the abundance of Tdh2-GFP protein. The median value of replicates is indicated with horizontal solid lines among groups of data points. \**p* < 0.05.

## Supplemental Tables

Table S1. Mutations in the evolved *TDH2-GFP-carrying* strain.

Table S2 Identities and frequencies of candidate mutations responsible for increased noise, and identified mutations in the ancestral-like and evolved-like F1 progeny pools.

Table S3. Plasmids and primers used in this study.

## References

1. Elowitz MB, Levine AJ, Siggia ED, Swain PS. Stochastic gene expression in a single cell. Science. 2002;297(5584):1183–6. Epub 2002/08/17. doi: 10.1126/science.1070919297/5584/1183 [pii]. PubMed PMID: 12183631.

2. Spudich JL, Koshland DE. Non-Genetic Individuality - Chance in Single Cell. Nature. 1976;262(5568):467–71. doi: DOI 10.1038/262467a0. PubMed PMID: WOS:A1976BZ68200026.

3. Suda T, Suda J, Ogawa M. Single-cell origin of mouse hemopoietic colonies expressing multiple lineages in variable combinations. Proc Natl Acad Sci U S A. 1983;80(21):6689–93. PubMed PMID: 6579554; PubMed Central PMCID: PMCPMC391236.

4. Losick R, Desplan C. Stochasticity and cell fate. Science. 2008;320(5872):65–8. Epub 2008/04/05. doi: 320/5872/65 [pii] 10.1126/science.1147888. PubMed PMID: 18388284; PubMed Central PMCID: PMC2605794.

5. Balazsi G, van Oudenaarden A, Collins JJ. Cellular decision making and biological noise: from microbes to mammals. Cell. 2011;144(6):910–25. Epub 2011/03/19. doi: S0092-8674(11)00069-9 [pii] 10.1016/j.cell.2011.01.030. PubMed PMID: 21414483; PubMed Central PMCID: PMC3068611.

6. Raj A, Rifkin SA, Andersen E, van Oudenaarden A. Variability in gene expression underlies incomplete penetrance. Nature. 2010;463(7283):913–8. doi: 10.1038/nature08781. PubMed PMID: 20164922; PubMed Central PMCID: PMCPMC2836165.

7. Burga A, Casanueva MO, Lehner B. Predicting mutation outcome from early stochastic variation in genetic interaction partners. Nature. 2011;480(7376):250–3. doi: 10.1038/nature10665. PubMed PMID: 22158248.

8. Acar M, Mettetal JT, van Oudenaarden A. Stochastic switching as a survival strategy in fluctuating environments. Nature Genetics. 2008;40(4):471–5. doi: 10.1038/ng.110. PubMed PMID: WOS:000254388100020.

9. Levy SF, Ziv N, Siegal ML. Bet hedging in yeast by heterogeneous, age-correlated expression of a stress protectant. PLoS Biol. 2012;10(5):e1001325. doi: 10.1371/journal.pbio.1001325. PubMed PMID: 22589700; PubMed Central PMCID: PMCPMC3348152.

10. de Jong IG, Haccou P, Kuipers OP. Bet hedging or not? A guide to proper classification of microbial survival strategies. Bioessays. 2011;33(3):215–23. Epub 2011/01/22. doi: 10.1002/bies.201000127. PubMed PMID: 21254151.

11. Raser JM, O’Shea EK. Control of stochasticity in eukaryotic gene expression. Science. 2004;304(5678):1811–4. Epub 2004/05/29. doi: 10.1126/science.10986411098641 [pii]. PubMed PMID: 15166317; PubMed Central PMCID: PMC1410811.

12. Bar-Even A, Paulsson J, Maheshri N, Carmi M, O’Shea E, Pilpel Y, et al. Noise in protein expression scales with natural protein abundance. Nat Genet. 2006;38(6):636–43. Epub 2006/05/23. doi: ng1807 [pii] 10.1038/ng1807. PubMed PMID: 16715097.

13. Taniguchi Y, Choi PJ, Li GW, Chen H, Babu M, Hearn J, et al. Quantifying E. coli proteome and transcriptome with single-molecule sensitivity in single cells. Science. 2010;329(5991):533–8. doi: 10.1126/science.1188308. PubMed PMID: 20671182; PubMed Central PMCID: PMCPMC2922915.

14. Newman JR, Ghaemmaghami S, Ihmels J, Breslow DK, Noble M, DeRisi JL, et al. Single-cell proteomic analysis of S. cerevisiae reveals the architecture of biological noise. Nature. 2006;441(7095):840–6. Epub 2006/05/16. doi: nature04785 [pii] 10.1038/nature04785. PubMed PMID: 16699522.

15. Barroso GV, Puzovic N, Dutheil JY. The Evolution of Gene-Specific Transcriptional Noise Is Driven by Selection at the Pathway Level. Genetics. 2018;208(1):173–89. doi: 10.1534/genetics.117.300457. PubMed PMID: WOS:000419356300012.

16. Bahar R, Hartmann CH, Rodriguez KA, Denny AD, Busuttil RA, Dolle ME, et al. Increased cell-to-cell variation in gene expression in ageing mouse heart. Nature. 2006;441(7096):1011–4. doi: 10.1038/nature04844. PubMed PMID: 16791200.

17. Hsieh YY, Hung PH, Leu JY. Hsp90 regulates nongenetic variation in response to environmental stress. Mol Cell. 2013;50(1):82–92. Epub 2013/02/26. doi: S1097-2765(13)00090-7 [pii] 10.1016/j.molcel.2013.01.026. PubMed PMID: 23434373.

18. Sanchez A, Choubey S, Kondev J. Regulation of noise in gene expression. Annu Rev Biophys. 2013;42:469–91. doi: 10.1146/annurev-biophys-083012-130401. PubMed PMID: 23527780.

19. Battich N, Stoeger T, Pelkmans L. Control of Transcript Variability in Single Mammalian Cells. Cell. 2015;163(7):1596–610. doi: 10.1016/j.cell.2015.11.018. PubMed PMID: 26687353.

20. Ansel J, Bottin H, Rodriguez-Beltran C, Damon C, Nagarajan M, Fehrmann S, et al. Cell-to-cell stochastic variation in gene expression is a complex genetic trait. PLoS Genet. 2008;4(4):e1000049. Epub 2008/04/12. doi: 10.1371/journal.pgen.1000049. PubMed PMID: 18404214; PubMed Central PMCID: PMC2289839.

21. Wu S, Li K, Li Y, Zhao T, Li T, Yang YF, et al. Independent regulation of gene expression level and noise by histone modifications. PLoS Comput Biol. 2017;13(6):e1005585. doi: 10.1371/journal.pcbi.1005585. PubMed PMID: 28665997; PubMed Central PMCID: PMCPMC5513504.

22. Freed NE, Silander OK, Stecher B, Bohm A, Hardt WD, Ackermann M. A simple screen to identify promoters conferring high levels of phenotypic noise. PLoS Genet. 2008;4(12):e1000307. doi: 10.1371/journal.pgen.1000307. PubMed PMID: 19096504; PubMed Central PMCID: PMCPMC2588653.

23. Carey JN, Mettert EL, Roggiani M, Myers KS, Kiley PJ, Goulian M. Regulated Stochasticity in a Bacterial Signaling Network Permits Tolerance to a Rapid Environmental Change. Cell. 2018;173(1):196–207 e14. doi: 10.1016/j.cell.2018.02.005. PubMed PMID: 29502970; PubMed Central PMCID: PMCPMC5866230.

24. Grant CM, Quinn KA, Dawes IW. Differential protein S-thiolation of glyceraldehyde-3-phosphate dehydrogenase isoenzymes influences sensitivity to oxidative stress. Mol Cell Biol. 1999;19(4):2650–6. PubMed PMID: 10082531; PubMed Central PMCID: PMCPMC84058.

25. Cohen D. Optimizing reproduction in a randomly varying environment. J Theor Biol. 1966;12(1):119–29. PubMed PMID: 6015423.

26. Venable DL. Bet hedging in a guild of desert annuals. Ecology. 2007;88(5):1086–90. doi: Doi 10.1890/06-1495. PubMed PMID: WOS:000246369900002.

27. Graham JK, Smith ML, Simons AM. Experimental evolution of bet hedging under manipulated environmental uncertainty in Neurospora crassa. Proc Biol Sci. 2014;281(1787). doi: 10.1098/rspb.2014.0706. PubMed PMID: 24870047; PubMed Central PMCID: PMCPMC4071552.

28. Stewart-Ornstein J, Weissman JS, El-Samad H. Cellular noise regulons underlie fluctuations in Saccharomyces cerevisiae. Mol Cell. 2012;45(4):483–93. doi: 10.1016/j.molcel.2011.11.035. PubMed PMID: 22365828; PubMed Central PMCID: PMCPMC3327736.

29. Konrad M. Analysis and in vivo disruption of the gene coding for adenylate kinase (ADK1) in the yeast Saccharomyces cerevisiae. J Biol Chem. 1988;263(36):19468–74. PubMed PMID: 2848829.

30. McNeil JB, McIntosh EM, Taylor BV, Zhang FR, Tang S, Bognar AL. Cloning and molecular characterization of three genes, including two genes encoding serine hydroxymethyltransferases, whose inactivation is required to render yeast auxotrophic for glycine. J Biol Chem. 1994;269(12):9155–65. PubMed PMID: 8132653.

31. Ahmad I, Rao DN. Functional analysis of conserved motifs in EcoP15I DNA methyltransferase. J Mol Biol. 1996;259(2):229–40. doi: 10.1006/jmbi.1996.0315. PubMed PMID: 8656425.

32. McBride AE, Weiss VH, Kim HK, Hogle JM, Silver PA. Analysis of the yeast arginine methyltransferase Hmt1p/Rmt1p and its in vivo function - Cofactor binding and substrate interactions. Journal of Biological Chemistry. 2000;275(5):3128–36. doi: DOI 10.1074/jbc.275.5.3128. PubMed PMID: WOS:000085146500018.

33. Gary JD, Lin WJ, Yang MC, Herschman HR, Clarke S. The predominant protein-arginine methyltransferase from Saccharomyces cerevisiae. J Biol Chem. 1996;271(21):12585–94. PubMed PMID: 8647869.

34. Li HT, Gong T, Zhou Z, Liu YT, Cao X, He Y, et al. Yeast Hmt1 catalyses asymmetric dimethylation of histone H3 arginine 2 in vitro. Biochem J. 2015;467(3):507–15. doi: 10.1042/BJ20141437. PubMed PMID: 25715670.

35. Jackson CA, Yadav N, Min S, Li J, Milliman EJ, Qu J, et al. Proteomic analysis of interactors for yeast protein arginine methyltransferase Hmt1 reveals novel substrate and insights into additional biological roles. Proteomics. 2012;12(22):3304–14. doi: 10.1002/pmic.201200132. PubMed PMID: WOS:000311616000005.

36. Lacoste N, Utley RT, Hunter JM, Poirier GG, Cote J. Disruptor of telomeric silencing-1 is a chromatin-specific histone H3 methyltransferase. Journal of Biological Chemistry. 2002;277(34):30421–4. doi: 10.1074/jbc.C200366200. PubMed PMID: WOS:000177579800004.

37. Yu MC, Lamming DW, Eskin JA, Sinclair DA, Silver PA. The role of protein arginine methylation in the formation of silent chromatin. Genes Dev. 2006;20(23):3249–54. doi: 10.1101/gad.1495206. PubMed PMID: 17158743; PubMed Central PMCID: PMCPMC1686602.

38. Wong CM, Tang HM, Kong KY, Wong GW, Qiu H, Jin DY, et al. Yeast arginine methyltransferase Hmt1p regulates transcription elongation and termination by methylating Npl3p. Nucleic Acids Res. 2010;38(7):2217–28. doi: 10.1093/nar/gkp1133. PubMed PMID: 20053728; PubMed Central PMCID: PMCPMC2853106.

39. Brandariz-Nunez A, Zeng F, Lam QN, Jin H. Sbp1 modulates the translation of Pab1 mRNA in a poly(A)- and RGG-dependent manner. RNA. 2018;24(1):43–55. doi: 10.1261/rna.062547.117. PubMed PMID: 28986506; PubMed Central PMCID: PMCPMC5733569.

40. Lipson RS, Webb KJ, Clarke SG. Rmt1 catalyzes zinc-finger independent arginine methylation of ribosomal protein Rps2 in Saccharomyces cerevisiae. Biochem Biophys Res Commun. 2010;391(4):1658–62. doi: 10.1016/j.bbrc.2009.12.112. PubMed PMID: 20035717; PubMed Central PMCID: PMCPMC2813213.

41. Bachand F, Silver PA. PRMT3 is a ribosomal protein methyltransferase that affects the cellular levels of ribosomal subunits. Embo Journal. 2004;23(13):2641–50. doi: 10.1038/sj.emboj.7600265. PubMed PMID: WOS:000223249300018.

42. Poornima G, Shah S, Vignesh V, Parker R, Rajyaguru PI. Arginine methylation promotes translation repression activity of eIF4G-binding protein, Scd6. Nucleic Acids Res. 2016;44(19):9358–68. doi: 10.1093/nar/gkw762. PubMed PMID: 27613419; PubMed Central PMCID: PMCPMC5100564.

43. Henry MF, Silver PA. A novel methyltransferase (Hmt1p) modifies poly(A)+-RNA-binding proteins. Mol Cell Biol. 1996;16(7):3668–78. PubMed PMID: 8668183; PubMed Central PMCID: PMCPMC231362.

44. Shen EC, Henry MF, Weiss VH, Valentini SR, Silver PA, Lee MS. Arginine methylation facilitates the nuclear export of hnRNP proteins. Genes Dev. 1998;12(5):679–91. PubMed PMID: 9499403; PubMed Central PMCID: PMCPMC316575.

45. Green DM, Marfatia KA, Crafton EB, Zhang X, Cheng XD, Corbett AH. Nab2p is required for poly(A) RNA export in Saccharomyces cerevisiae and is regulated by arginine methylation via Hmt1p. Journal of Biological Chemistry. 2002;277(10):7752–60. doi: DOI 10.1074/jbc.M110053200. PubMed PMID: WOS:000174268000018.

46. Chen YC, Milliman EJ, Goulet I, Cote J, Jackson CA, Vollbracht JA, et al. Protein arginine methylation facilitates cotranscriptional recruitment of pre-mRNA splicing factors. Mol Cell Biol. 2010;30(21):5245–56. doi: 10.1128/MCB.00359-10. PubMed PMID: 20823272; PubMed Central PMCID: PMCPMC2953043.

47. Laurent BC, Treitel MA, Carlson M. Functional interdependence of the yeast SNF2, SNF5, and SNF6 proteins in transcriptional activation. Proc Natl Acad Sci U S A. 1991;88(7):2687–91. PubMed PMID: 1901413; PubMed Central PMCID: PMCPMC51303.

48. Cairns BR, Kim YJ, Sayre MH, Laurent BC, Kornberg RD. A multisubunit complex containing the SWI1/ADR6, SWI2/SNF2, SWI3, SNF5, and SNF6 gene products isolated from yeast. Proc Natl Acad Sci U S A. 1994;91(5):1950–4. PubMed PMID: 8127913; PubMed Central PMCID: PMCPMC43282.

49. Ansari SA, Paul E, Sommer S, Lieleg C, He QY, Daly AZ, et al. Mediator, TATA-binding Protein, and RNA Polymerase II Contribute to Low Histone Occupancy at Active Gene Promoters in Yeast. Journal of Biological Chemistry. 2014;289(21):14981–95. doi: 10.1074/jbc.M113.529354. PubMed PMID: WOS:000337248100050.

50. Alejandro-Osorio AL, Huebert DJ, Porcaro DT, Sonntag ME, Nillasithanukroh S, Lwill J, et al. The histone deacetylase Rpd3p is required for transient changes in genomic expression in response to stress. Genome Biology. 2009;10(5). doi: ARTN R57 10.1186/gb-2009-10-5-r57. PubMed PMID: WOS:000267604200016.

51. Worley J, Luo X, Capaldi AP. Inositol pyrophosphates regulate cell growth and the environmental stress response by activating the HDAC Rpd3L. Cell Rep. 2013;3(5):1476–82. doi: 10.1016/j.celrep.2013.03.043. PubMed PMID: 23643537; PubMed Central PMCID: PMCPMC3672359.

52. Taylor JW, Berbee ML. Dating divergences in the Fungal Tree of Life: review and new analyses. Mycologia. 2006;98(6):838–49. Epub 2007/05/10. PubMed PMID: 17486961.

53. Booth IR. Stress and the single cell: intrapopulation diversity is a mechanism to ensure survival upon exposure to stress. Int J Food Microbiol. 2002;78(1-2):19–30. Epub 2002/09/12. PubMed PMID: 12222634.

54. Fraser D, Kaern M. A chance at survival: gene expression noise and phenotypic diversification strategies. Mol Microbiol. 2009;71(6):1333–40. Epub 2009/02/18. doi: MMI6605 [pii] 10.1111/j.1365-2958.2009.06605.x. PubMed PMID: 19220745.

55. Veening JW, Smits WK, Kuipers OP. Bistability, epigenetics, and bet-hedging in bacteria. Annu Rev Microbiol. 2008;62:193–210. Epub 2008/06/10. doi: 10.1146/annurev.micro.62.081307.163002. PubMed PMID: 18537474.

56. Stark C, Breitkreutz BJ, Reguly T, Boucher L, Breitkreutz A, Tyers M. BioGRID: a general repository for interaction datasets. Nucleic Acids Res. 2006;34(Database issue):D535–9. Epub 2005/12/31. doi: 34/suppl_1/D535 [pii] 10.1093/nar/gkj109. PubMed PMID: 16381927; PubMed Central PMCID: PMC1347471.

57. Gopinath RK, You ST, Chien KY, Swamy KB, Yu JS, Schuyler SC, et al. The Hsp90-dependent proteome is conserved and enriched for hub proteins with high levels of protein-protein connectivity. Genome Biol Evol. 2014;6(10):2851–65. Epub 2014/10/16. doi: evu226 [pii] 10.1093/gbe/evu226. PubMed PMID: 25316598; PubMed Central PMCID: PMC4224352.

58. Albert R, Jeong H, Barabasi AL. Error and attack tolerance of complex networks. Nature. 2000;406(6794):378–82. Epub 2000/08/10. doi: 10.1038/35019019. PubMed PMID: 10935628.

59. Bergman A, Siegal ML. Evolutionary capacitance as a general feature of complex gene networks. Nature. 2003;424(6948):549–52. doi: Doi 10.1038/Nature01834. PubMed PMID: ISI:000184454700042.

60. Levy SF, Siegal ML. Network hubs buffer environmental variation in Saccharomyces cerevisiae. PLoS Biol. 2008;6(11):e264. Epub 2008/11/07. doi: 08-PLBI-RA-0960 [pii] 10.1371/journal.pbio.0060264. PubMed PMID: 18986213; PubMed Central PMCID: PMC2577700.

61. Synetos D, Frantziou CP, Alksne LE. Mutations in yeast ribosomal proteins S28 and S4 affect the accuracy of translation and alter the sensitivity of the ribosomes to paromomycin. Biochim Biophys Acta. 1996;1309(1-2):156–66. PubMed PMID: 8950190.

62. Rother S, Strasser K. The RNA polymerase II CTD kinase Ctk1 functions in translation elongation. Genes Dev. 2007;21(11):1409–21. doi: 10.1101/gad.428407. PubMed PMID: 17545469; PubMed Central PMCID: PMCPMC1877752.

63. Blanc RS, Richard S. Arginine Methylation: The Coming of Age. Mol Cell. 2017;65(1):8–24. doi: 10.1016/j.molcel.2016.11.003. PubMed PMID: 28061334.

64. Low JKK, Wilkins MR. Protein arginine methylation in Saccharomyces cerevisiae. Febs Journal. 2012;279(24):4423–43. doi: 10.1111/febs.12039. PubMed PMID: WOS:000312224600001.

65. Bedford MT, Clarke SG. Protein Arginine Methylation in Mammals: Who, What, and Why. Molecular Cell. 2009;33(1):1–13. doi: 10.1016/j.molcel.2008.12.013. PubMed PMID: WOS:000262624700001.

66. Xu W, Cho H, Kadam S, Banayo EM, Anderson S, Yates JR, 3rd, et al. A methylation-mediator complex in hormone signaling. Genes Dev. 2004;18(2):144–56. doi: 10.1101/gad.1141704. PubMed PMID: 14729568; PubMed Central PMCID: PMCPMC324421.

67. Ito H, Fukuda Y, Murata K, Kimura A. Transformation of intact yeast cells treated with alkali cations. J Bacteriol. 1983;153(1):163–8. PubMed PMID: 6336730.

68. Hsu PC, Yang CY, Lan CY. Candida albicans Hap43 Is a Repressor Induced under Low-Iron Conditions and Is Essential for Iron-Responsive Transcriptional Regulation and Virulence. Eukaryotic Cell. 2011;10(2):207–25. doi: 10.1128/Ec.00158-10. PubMed PMID: WOS:000286981300007.

69. Huh WK, Falvo JV, Gerke LC, Carroll AS, Howson RW, Weissman JS, et al. Global analysis of protein localization in budding yeast. Nature. 2003;425(6959):686–91. Epub 2003/10/17. doi: 10.1038/nature02026 nature02026 [pii]. PubMed PMID: 14562095.

70. Subach OM, Cranfill PJ, Davidson MW, Verkhusha VV. An enhanced monomeric blue fluorescent protein with the high chemical stability of the chromophore. PLoS One. 2011;6(12):e28674. doi: 10.1371/journal.pone.0028674. PubMed PMID: 22174863; PubMed Central PMCID: PMCPMC3234270.

71. Metzger BP, Yuan DC, Gruber JD, Duveau F, Wittkopp PJ. Selection on noise constrains variation in a eukaryotic promoter. Nature. 2015;521(7552):344–7. doi: 10.1038/nature14244. PubMed PMID: 25778704; PubMed Central PMCID: PMCPMC4455047.

72. Nagai T, Ibata K, Park ES, Kubota M, Mikoshiba K, Miyawaki A. A variant of yellow fluorescent protein with fast and efficient maturation for cell-biological applications. Nat Biotechnol. 2002;20(1):87–90. doi: 10.1038/nbt0102-87. PubMed PMID: 11753368.

73. Winzeler EA, Shoemaker DD, Astromoff A, Liang H, Anderson K, Andre B, et al. Functional characterization of the S-cerevisiae genome by gene deletion and parallel analysis. Science. 1999;285(5429):901–6. doi: DOI 10.1126/science.285.5429.901. PubMed PMID: WOS:000081860900053.

74. Breslow DK, Cameron DM, Collins SR, Schuldiner M, Stewart-Ornstein J, Newman HW, et al. A comprehensive strategy enabling high-resolution functional analysis of the yeast genome. Nat Methods. 2008;5(8):711–8. doi: 10.1038/nmeth.1234. PubMed PMID: 18622397; PubMed Central PMCID: PMCPMC2756093.

75. DiCarlo JE, Norville JE, Mali P, Rios X, Aach J, Church GM. Genome engineering in Saccharomyces cerevisiae using CRISPR-Cas systems. Nucleic Acids Research. 2013;41(7):4336–43. doi: 10.1093/nar/gkt135. PubMed PMID: WOS:000318167900046.

76. Horwitz AA, Walter JM, Schubert MG, Kung SH, Hawkins K, Platt DM, et al. Efficient Multiplexed Integration of Synergistic Alleles and Metabolic Pathways in Yeasts via CRISPR-Cas. Cell Syst. 2015;1(1):88–96. doi: 10.1016/j.cels.2015.02.001. PubMed PMID: 27135688.

77. Liu J, Huang S, Sun M, Liu S, Liu Y, Wang W, et al. An improved allele-specific PCR primer design method for SNP marker analysis and its application. Plant Methods. 2012;8(1):34. doi: 10.1186/1746-4811-8-34. PubMed PMID: 22920499; PubMed Central PMCID: PMCPMC3495711.

78. Lenski RE, Rose MR, Simpson SC, Tadler SC. Long-Term Experimental Evolution in Escherichia-Coli.1. Adaptation and Divergence during 2,000 Generations. American Naturalist. 1991;138(6):1315–41. doi: Doi 10.1086/285289. PubMed PMID: WOS:A1991HB55000001.

79. Winston F. EMS and UV mutagenesis in yeast. Curr Protoc Mol Biol. 2008;Chapter 13:Unit 13 3B. doi: 10.1002/0471142727.mb1303bs82. PubMed PMID: 18425760.

80. Ozbudak EM, Thattai M, Kurtser I, Grossman AD, van Oudenaarden A. Regulation of noise in the expression of a single gene. Nat Genet. 2002;31(1):69–73. Epub 2002/04/23. doi: 10.1038/ng869 ng869 [pii]. PubMed PMID: 11967532.

81. Teng X, Cheng WC, Qi B, Yu TX, Ramachandran K, Boersma MD, et al. Gene-dependent cell death in yeast. Cell Death Dis. 2011;2:e188. doi: 10.1038/cddis.2011.72. PubMed PMID: 21814286; PubMed Central PMCID: PMCPMC3181418.

82. Lee HY, Cheng KY, Chao JC, Leu JY. Differentiated cytoplasmic granule formation in quiescent and non-quiescent cells upon chronological aging. Microb Cell. 2016;3(3):109–19. PubMed PMID: WOS:000373126700004.

83. Kushnirov VV. Rapid and reliable protein extraction from yeast. Yeast. 2000;16(9):857–60. Epub 2000/06/22. doi: 10.1002/1097-0061(20000630)16:9<857::AID-YEA561>3.0.CO;2-B [pii] 10.1002/1097-0061(20000630)16:9<857::AID-YEA561>3.0.CO;2-B. PubMed PMID: 10861908.

84. Takagi H, Abe A, Yoshida K, Kosugi S, Natsume S, Mitsuoka C, et al. QTL-seq: rapid mapping of quantitative trait loci in rice by whole genome resequencing of DNA from two bulked populations. Plant J. 2013;74(1):174–83. doi: 10.1111/tpj.12105. PubMed PMID: 23289725.

85. Lee W, Tillo D, Bray N, Morse RH, Davis RW, Hughes TR, et al. A high-resolution atlas of nucleosome occupancy in yeast. Nat Genet. 2007;39(10):1235–44. doi: 10.1038/ng2117. PubMed PMID: 17873876.

86. Keren L, van Dijk D, Weingarten-Gabbay S, Davidi D, Jona G, Weinberger A, et al. Noise in gene expression is coupled to growth rate. Genome Research. 2015;25(12):1893–902. doi: 10.1101/gr.191635.115. PubMed PMID: WOS:000365830400011.

